# Experimental type 1 diabetes metabolically rejuvenates CD8^+^ T cells for improved control of tumor growth through an IGF1-IGF1R axis

**DOI:** 10.1101/2024.04.04.588206

**Authors:** Anirban Sarkar, Sukanya Dhar, Saurav Bera, Mohona Chakravarti, Ayushi Verma, Parash Prasad, Jasmine Sultana, Juhina Das, Akata Saha, Avishek Bhuniya, Ipsita Guha, Shayani Dasgupta, Sib Sankar Roy, Saptak Banerjee, Subir Roy, Debarati Bhar, Walter J. Storkus, Rathindranath Baral, Dipak Datta, Anamika Bose

## Abstract

Epidemiological studies suggest that patients with pre-existing type 1 diabetes (T1D) have a decreased risk of developing melanoma, prostate cancer, and breast cancer, although the underlying mechanism remains to be elucidated. In translational modelling, we observed that streptozotocin (STZ) induced T1D mice exhibited restricted melanoma and carcinoma (mammary, lung and colon) growth in association with extended overall survival. Tumor-infiltrating CD8^+^ T cells were found to be responsible for tumor growth restriction. Tumor infiltrating CD8^+^ T cells but not tumor cells themselves exhibited higher glycolytic and cytotoxic activities in T1D hosts. Such improved anti-tumor T cell function was linked to selective upregulated expression of insulin-like growth factor 1, insulin-like growth factor 1 receptor, and phospho-mTOR in CD8^+^ T cells in the TME. T1D patient derived CD8^+^ T cells displayed superior activation *in vitro* after tumor antigen stimulation vs. non-diabetic CD8^+^ T cells. Activation of T1D patient derived CD8^+^ T cells was sensitive to targeted antagonism of IGF1R and mTOR, supporting the operational involvement of the IGF1R-mTOR signaling axis. Our results suggest that selective activation of the intrinsic IGF1R-mTOR signaling axis in CD8^+^ T cells represents a preferred endpoint to achieving more effective immunotherapy outcomes and improved cancer patient management.

**Significance:** Experimental type 1 diabetes decelerates tumor growth through metabolic activation of cytotoxic T cells dependent on an IGF1R-mTOR signaling pathway. CD8^+^IGF1R^+^IGF1^+^ T cells play a crucial role in T1D dependent tumor control.

## Introduction

The incidence of diabetes is increasing globally and represents a leading cause of death annually. Type 1 diabetes (T1D) or insulin-dependent diabetes mellitus (IDDM), a major form of diabetes, is an autoimmune disease characterized by hyperglycemia and insulin deficiency due to CD8^+^ T cell-targeted loss of pancreatic β cells (1). Preclinical and clinical evidence in T1D models/patients suggests that CD8^+^ T cell infiltration not only occurs in pancreatic tissue, but also in other peripheral organs, including the liver, kidney, brain, and eye (2,3). Such inflammatory infiltrates cause significant damage to these organs, resulting in pathologic hepatopathy, nephropathy, neuropathy, and retinopathy. Untreated T1D can lead to death. Since stem cell therapy to restore pancreatic beta cells and immune cell-based therapies designed to therapeutically treat T1D remain in their developmental stages, administration of insulin represents the sole life sustaining therapy for patients with T1D (4). Provision of insulin restores patient blood glucose levels to normal and serves to regulate systemic inflammation (5). Since the incidence of T1D is increasing annually among children, there is an increasing need for further research to study its impact on inflammatory diseases, including cancer (6). Indeed, the underlying mechanism(s) by which T1D may influence cancer progression remains largely unknown.

CD8^+^ T cells play a major role in preventing and controlling cancer development and progression, with cytotoxic T lymphocytes (CTLs) capable of directly killing tumor cells (7,8). Limited exploration has been done so far to determine the possible influence of pre-existing T1D or T1D associated activation of CTL in the cancer setting. Meta-analysis, using PubMed and EMBASE observational studies, suggests an association between T1D and increased risk for developing cancers of the stomach, lung, pancreas, liver, ovary, and kidney, but a decreased risk for melanoma, prostate, and breast cancer (9). We hypothesized that in T1D with at least certain forms of cancer, that CD8^+^ T cells may infiltrate tumors and mediate superior anti-tumor activity that is beneficial for patients.

A retrospective study by Carstensen et al., 2016 of 9000 cancer patients with T1D spanning 5 countries suggested a reduced risk of melanoma, breast carcinoma or prostate cancer, but an increased risk for patients developing colon, esophageal, liver or stomach cancer. This observation indicates contextual association of T1D with varied forms of cancers. A translational study by Heuson and Legros, 1972 investigating rat mammary carcinomas in an alloxan-induced T1D model reported tumor regression in 90% of diabetic rats; however, the underlying mechanism for host protection was largely unexplored. Based on these unresolved findings, in the present study we examined the influence of streptozotocin-induced pre-existing T1D on cancer progression and associated immune alterations in a range of murine melanoma and carcinoma (i.e., breast, colon and lung) models. The streptozotocin induce T1D model was employed as this system is widely used in the field of endocrinology research (18,19). We found that the presence of T1D in the host yields an immunologically “hot” TME in which tumor growth is effectively controlled by anti-tumor CTL. Our results further provide mechanistic insight into T1D-induced metabolic reprogramming of CTL via the IGF1R-mTOR signaling axis which may be actionable for the design and implementation of improved immunotherapies for cancer patients.

## Results

### 1. Pre-existing T1D is associated with reduced growth of transplantable tumors resulting in extended overall survival

To examine the influence of pre-existing T1D on tumor progression, tumor growth was studied in a low dose streptozotocin (STZ) induced murine T1D model. (experimental design is presented in **Figure 1A**). Prior to tumor inoculation, T1D induced mice were screened for blood glucose levels, with mice exhibiting blood glucose levels between 250-350 mg/dl selected as recipients for tumor inoculation **(Figure S1A)**. Such screening was done to eliminate strain specific STZ sensitivity, with control groups of mice displaying a mean blood glucose level of 100 mg/dl. Insulin administration significantly reduced blood glucose concentrations in C57BL/6J and BALB/c mice. Serum insulin levels in both strains of mice was significantly reduced following STZ treatment **(Figure S1B)**. STZ induced T1D mice showed significantly slowed growth of B16F10, 4T1, LLC and CT26 tumors as monitored on days 21 and 28, unless they received treatment with insulin (**Figure 1B and 1C**). STZ induced T1D mice bearing B16F10, 4T1, LLC or CT26 tumors also displayed a significant survival benefit when compared to untreated (control) and STZ+insulin treated groups of mice (**Figure 1D**).

**Figure 1.**
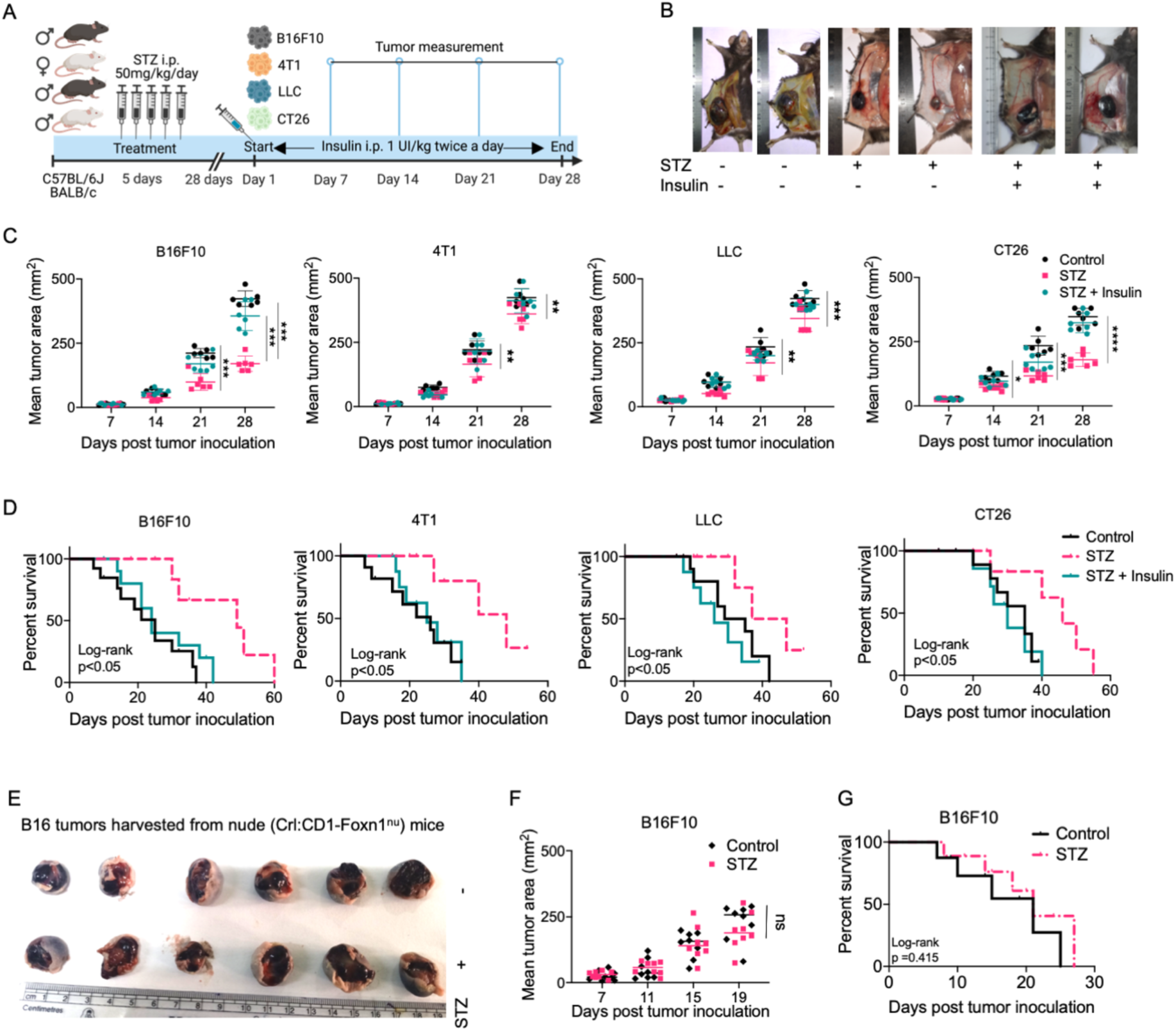
Pre-existing T1D is associated with reduced growth of transplantable tumors resulting in extended overall survival. **(A)** Schematic representation of experimental design. Three hosts were investigated:, untreated or control, STZ induced T1D and STZ induced T1D treated with insulin (n=6 male or female C57BL/6 or BALB/c mice/cohort). STZ was administered i.p. for five consecutive days and animals then left untreated for 28 days to allow for T1D to develop. On day 1, each of the 3 host groups were injected with tumors (s.c. B16F10 melanoma cells or LLC lung carcinoma cells in male C57BL/6 mice, or 4T1 breast carcinoma cells into the mammary fat pad in female BALB/c mice or s.c. CT-26 colon carcinoma cells in female BALB/c mice). Insulin was then administered i.p. to the STZ+insulin host groups twice a day throughout the course of the experiments. (**B)** Representative photographs of B16F10 tumor bearing C57BL/6J mice with scale rulers collected at the end of the experiment (day 28). (**C)** Mean tumor area (mm^2^) of B16F10, 4T1, LLC and CT26 tumor bearing mice with control, STZ treated and STZ+insulin treated (n=6) groups. Tumor areas were determined in each model on days 7, 14, 21 and 28. (**D)** Kaplan-Meier survival curve of B16F10, 4T1, LLC and CT26 tumor bearing mice in control, STZ induced T1D and STZ induced T1D + insulin treated hosts. Survival curves were plotted and intergroup differences analyzed using a log-rank test. **(E)** Representative photographs (with scale rulers) of B16F10 tumors harvested from untreated and STZ induced T1D nude (Crl:CD1-Foxn1^nu^) mice (day 19). **(F)** Mean B16F10 tumor area (mm^2^) in nude mice on days 7, 11, 15 and 19. **(G)** Kaplan-Meier survival curve of B16F10 melanoma bearing nude mice with intergroup differences analysed using a log rank test. Two-way ANOVA, followed by Tukey’s multiple comparisons test was performed for C and F. (*P ≤ 0.05, **P ≤ 0.01, ***P ≤ 0.001, ****P ≤ 0.0001).

To determine whether slowed tumor growth in T1D mice was mediated by the adaptive immune system, we performed analogous experiments in nude (Crl:CD1-Foxn^nu^) recipient mice. Nude mice showed significantly elevated serum glucose and reduced serum insulin levels following STZ treatment prior to tumor inoculation **(Figure S1E and S1F)**. Nude mice treated with STZ failed to restrict B16F10 tumor growth vs. control mice (**Figure 1E and 1F**), with no significant survival benefit observed vs. tumor-bearing non-diabetic nude mice (**Figure 1G**). These results support (adaptive) immune system-mediated control of tumor growth under T1D physiologic conditions that is mitigated upon insulin administration.

Among serum cytokines evaluated prior to tumor inoculation in STZ-induced T1D mice, we observed a significant increase in levels of IL-6 in both mouse strains, which was normalized by insulin treatment **(Figure S1D)**. In contrast, serum levels of IL-17, IFNγ, TNFα and IL-10 remained comparable across the untreated (control), STZ treated and STZ+insulin treated groups. Histologic and immunohistochemical analyses of the pancreas revealed reduced pancreatic islet content following STZ treatment in C57BL/6 mice **(Figure S1G and S1H)**, with livers in STZ treated mice exhibiting enlarged hepatocytes **(Figure S1I)**.

### 2. Existing T1D is associated with the increased recruitment and anti-tumor effector function of CD8^+^ T cells

Given the involvement of adaptive immunity in T1D-associated tumor growth control, we next analyzed T cell content in the TME. CD8^+^ T cells were found to be significantly upregulated amongst the tumor infiltrating lymphocyte (TIL) population in STZ induced T1D mice bearing B16F10, 4T1, LLC or CT26 tumors when compared to non-diabetic tumor control cohorts, which was mitigated by treatment with insulin (**Figure 2A**). CD8^+^CD28^+^ TILs isolated from B16F10, 4T1, LLC or CT26 tumor bearing STZ induced T1D hosts displayed significantly upregulated expression of CD69 when compared to their non-diabetic tumor control counterparts. Insulin treatment (STZ+insulin) reduced TIL expression of CD69 in the B16F10 tumor model but not in the 4T1, LLC or CT26 tumor models (**Figure 2B**). Remarkably, CD8^+^ TILs isolated from STZ treated B16F10, 4T1, LLC and CT26 tumor bearing T1D mice expressed enhances levels of CXCR3 (**Figure 2C**), granzyme B (GZMB) **(Figure S2A and S2B)** and interferon γ (IFNγ) **(Figure S2C)** when compared to untreated control TILs. Insulin treatment of these STZ induced T1D tumor models normalized the population of CD8^+^CXCR3^+^ TIL in the 4T1 and LLC bearing mice, but not in B16F10 or CT26 bearing mice. Furthermore, insulin treatment normalized expression of GZMB and IFNγ in TILs isolated from all four T1D tumor models evaluated. Tumors in T1D mice exhibited significantly higher CD8^+^CD103^+^CD39^+^ TIL content compared to control groups, which was reversible upon treatment with insulin **(Figure S3A)**. Higher frequencies of CD8^+^CD69^+^GZMB^+^ T cells were detected in STZ treated tumor draining lymph nodes (TDLNs), but not in the spleens of B16F10, 4T1, LLC or CT26 tumor bearing T1D mice, which were counteracted by insulin treatment **(Figure S3B and S3C)**.

**Figure 2.**
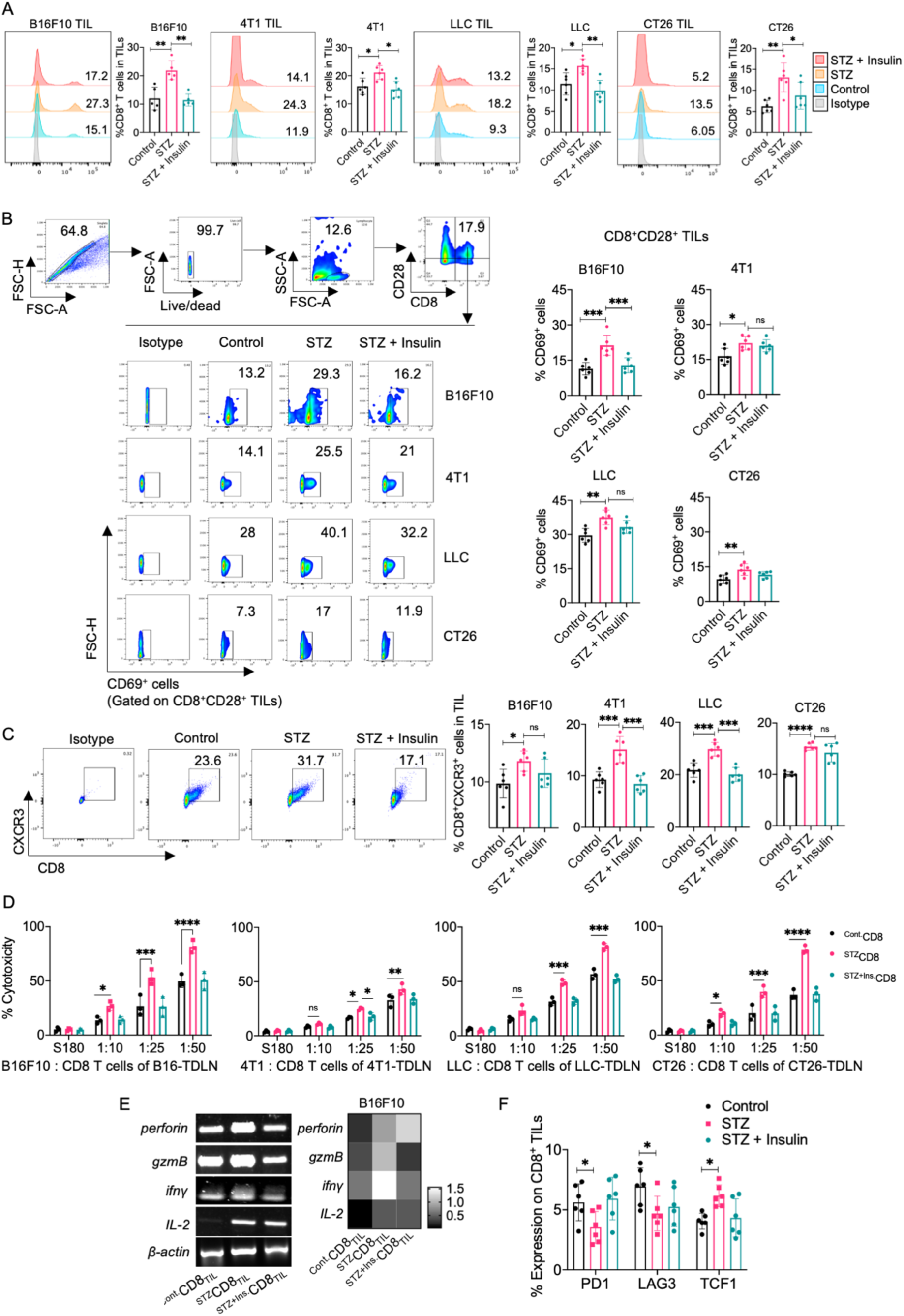
Existing T1D is associated with the increased recruitment and anti-tumor effector function of CD8^+^ T cells. **(A)** Histogram and graphical quantitation of CD8^+^ TIL contents in B16F10, 4T1, LLC and CT26 tumors in each recipient host evaluated (n=6 mice/group). (**B)** Gating strategy for flow analysis of CD8^+^, CD28^+^ and CD69^+^ cell events along with a representative flow cytometry data set representative of TIL isolated from control, STZ treated and STZ+insulin treated hosts (n=6 mice/group) bearing B16F10, 4T1, LLC or CT26 tumors. Graphs depict the percentage of CD69^+^ cells amongst CD8^+^CD28^+^ TILs. **(C)** Representative gating and graphical data reporting the percentage of CD8^+^CXCR3^+^ TIL in B16F10, 4T1, LLC and CT26 tumors in control, STZ treated and STZ+insulin treated hosts (n=6 mice/group). (**D)** Graphs depicting the percentage cytotoxicity of model-matched tumor cells mediated by CD8^+^ TILs isolated from B16F10, 4T1, LLC and CT26 tumors (LDH release assay). LDH release of control, STZ treated and STZ+insulin treated (n=3, repeated twice) groups were compared. Cancer cell : CD8^+^ T cell ratios of 1:10, 1:25 and 1:50 were used. S180 sarcoma cells were used as unrelated control target. **(E)** Heatmap depicts relative expression of *perforin, granzyme B, ifnγ and il-2* genes in ^Cont.^CD8_TIL_, ^STZ^CD8_TIL_ and ^STZ+ins.^CD8_TIL_ cells sorted from B16F10 tumor-bearing mice. **(F)** Percentage of CD8^+^ TILs isolated from B16F10 tumors that co-express PD1, LAG3 or TCF1 (n=6 mice/group). One way ANOVA followed by Tukey’s multiple comparisons test was performed to determine the significance of inter-group differences. (*P ≤ 0.05, **P ≤ 0.01, ***P ≤ 0.001, ****P ≤ 0.0001)

To assess model-associated changes in CD8^+^ T cell-mediated cytotoxicity, we employed CD8-MACS to isolate CD8^+^ T cells from TDLNs harvested from B16F10, 4T1, LLC and CT26 tumor bearing mice (i.e., control, STZ induced, and STZ induced +insulin treated). Sorted CD8^+^ T cells were cultured *in vitro* with isologous tumor cells at different effector : target (E : T) ratios, with lactate dehydrogenase (LDH) release measured as an effector cell readout. CD8^+^ T cells from B16F10, 4T1, LLC and CT26-bearing T1D mice mediated significantly enhanced cytotoxicity against their model relevant (but not irrelevant) tumor cell targets, which was substantially reduced in CD8^+^ T cells isolated from T1D + insulin treated tumor-bearing mice (**Figure 2D**). Similar differences for CD8^+^ TIL expression of *perforin*, *grB*, *IFNg* and *il-2* in B16F10 tumor bearing T1D vs. T1D + insulin treated mice (**Figure 2E**). In extended analyses, we observed that CD8^+^ TIL isolated from melanoma-bearing STZ induced T1D (vs. control) mice expressed reduced levels of the PD1 and LAG3 checkpoint molecules, but elevated expression of TCF1 (**Figure 2F**). Analyses of immune-suppressor cells in the TME indicated decreased frequencies of CD4^+^FoxP3^+^ Tregs, CD11b^+^F4/80^+^ TAMs and CD11b^+^Gr1^+^ MDSCs in STZ induced mice bearing B16F10 tumors, with unaltered content of CD11c^+^MHC-I^+^ DCs when compared to control tumors **(Figure S4A, Figure S4B)**. These cumulative results suggest pre-existing T1D host facilitates an immunologically “hot” TME based on preferential recruitment/maintenance of pro-inflammatory immune cell populations in melanoma and carcinoma tumors and exclusion/removal of regulatory cell populations in melanoma tumors which is leveraged by the absence or presence of insulin.

### 3. T1D associated tumor growth control is CD8^+^ T cell dependent and may be adoptively transferred

Given the direct involvement of the immune system, particularly CD8^+^ T cells, in diabetes (20–22) as well as in cancer (23,24), the influence of T1D on CD8^+^ T effector cells in our tumor models was next assessed. As described above, STZ administration and maintained for 4 weeks to develop T1D in both C57BL/6J (n=6) and nude mice (n=6), with a group of nondiabetic C57BL/6J (n=6) and nude (n=6) mice serving as controls. B16F10 tumors were injected s.c. in mice. After palpable tumor formation, CD8^+^ T cells were harvested from the TDLN of control as well as STZ-induced B16F10 bearing mice by MACS. Isolated CD8^+^ T cells (1x10^6^ cells/mice) from control or STZ-induced T1D mice were adoptively transferred into control or STZ induced nude mice bearing B16F10 tumors by tail vein injection (**Figure 3A**). We observed that although tumor growth was reduced upon adoptive transfer of CD8^+^ T cells from either source (vs. no transfer) in both the non-diabetic and T1D nude melanoma models, that adoptive T cell therapy implementing CD8^+^ T cells isolated from T1D tumor bearing mice (i.e., ^STZ^CD8^+^ T cells) mediated significantly greater tumor growth control (p<0.05) in T1D hosts when compared to ACT using CD8^+^ T cells isolated from non-diabetic tumor bearing mice (**Figure 3B and 3C**).

**Figure 3.**
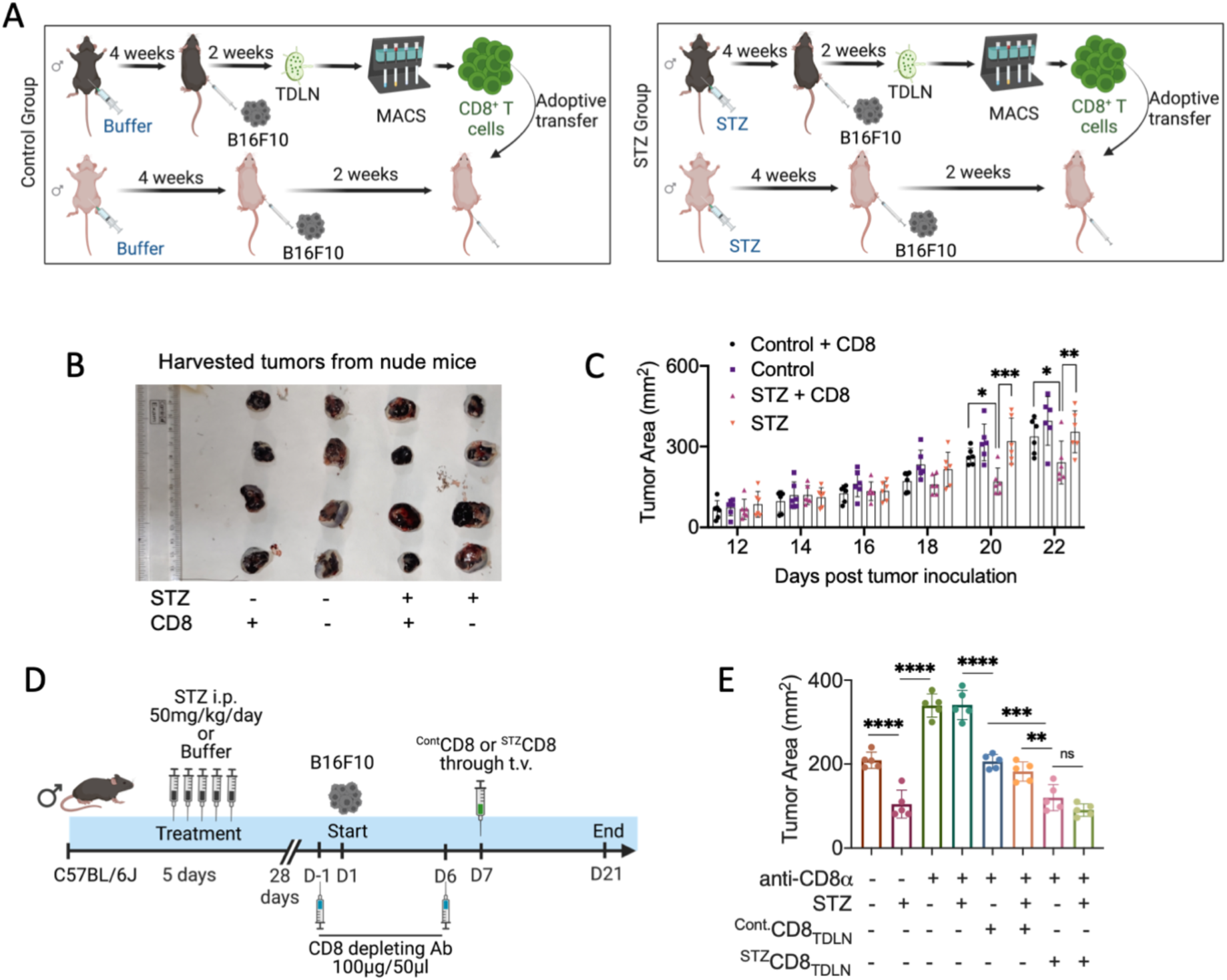
T1D associated tumor growth control is CD8^+^ T cell dependent and may be adoptively transferred. **(A)** Schematic overview of experimental design. C57BL/6J (black) and nude (pink) mice were divided into two groups. One group received streptozotocin (STZ) i.p. (50mg/kg) for five consecutive days, while the other group received only buffer (Sodium citrate, pH 4.5), with animals subsequently maintained for four weeks to allow for diabetes induction. After four weeks, STZ treated mice with random blood sugar values in the range of 250-350 mg/dl were selected for experimental use. B16F10 cells (5×10^5^ cells) were injected s.c. into C57BL/6 mice and allowed to establish palpable tumors. Two weeks after tumor injection, animals were euthanized and CD8^+^ T cells (^STZ^CD8_TDLN_ or ^Cont.^CD8_TDLN_) were isolated from tumor draining lymph nodes (TDLNs) by magnetic assisted cell sorting (MACS). MACS sorted ^STZ^CD8_TDLN_ or ^Cont.^CD8_TDLN_ (2x10^5^ cells/mice) or PBS (as a control) were adoptively transferred via tail vein injection into nude mice bearing established B16F10 melanomas. Tumor growth was monitored through day 22. (**B)** Representative photographs of B16F10 tumors in nude mice harvested 22 days after tumor inoculation. (**C)** Mean tumor area in each of the different experimental groups (n=6) is plotted graphically over time. **(D)** Control or STZ induced T1D C57BL/6J mice (n=5 mice/group) were injected s.c. with B6F10 melanoma cells on study day 1. Anti CD8a antibody (100μg/50μl PBS was injected i.p. on days -1 and 6 in the indicated groups. On day 7, cohorts of control or STZ induced T1D mice bearing B16F10 melanomas received adoptive transfer of ^STZ^CD8_TDLN_ or ^Cont.^CD8_TDLN_ T cells (2 x10^5^ cells/mouse) via tail vein injection as indicated. Tumor area were measured on day 28 post-tumor inoculation. **(E)** Mean B16F10 tumor area is reported for each experimental cohort from (D) Two way ANOVA followed by Tukey’s multiple comparisons test was performed to calculate the significance or inter-cohort differences. (*P ≤ 0.05, **P ≤ 0.01, ***P ≤ 0.001, ****P ≤ 0.0001)

For further understanding on the role of CD8^+^ T cells in controlling tumor growth in STZ induced T1D mice, systemic CD8^+^ T cell depletion was performed (**Figure 3D**). C57BL/6J mice were divided into eight groups (n=5) with half receiving STZ treatment (50 mg/kg for consecutive 5 days) and the other four groups left untreated. Depleting anti-CD8a antibody (100μg/50μl) was injected i.p on days -1 and 6, into 3 of the untreated and 3 of the STZ induced groups. B16F10 tumors were injected s.c. on day 1 into all animals. Following palpable tumor formation on day 7, CD8^+^ T cells were isolated from TDLN of untreated (^Cont.^CD8_TDLN_) or STZ treated (^STZ^CD8_TDLN_) tumor bearing mice. One untreated and one STZ induced recipient group received adoptive T cell therapy implementing ^Cont.^CD8_TDLN_ or ^STZ^CD8_TDLN_. CD8^+^ T cell depletion in T1D but not recipient tumor bearing mice ablated tumor growth control vs. non-diabetic CD8^+^ T cell depleted tumor bearing hosts (**Figure 3E**). Adoptive CD8^+^ T cell therapy resulted in tumor growth restriction in both non-diabetic and STZ treated groups receiving either ^Cont^CD8_TDLN_ or ^STZ^CD8_TDLN_ T cells, compared to their respective control groups received no adoptive T cell therapy. However, non-diabetic and STZ treated groups that received ^STZ^CD8_TDLN_ therapy showed more prominent tumor growth restriction than non-diabetic and STZ treated groups that received ^Cont^CD8_TDLN_ therapy (p<0.01). Collectively, these results confirmed the involvement of effector CD8^+^ T cells in restricting tumor growth in T1D tumor-bearing hosts.

### 4. T1D metabolically reprograms CD8^+^ T lymphocytes in support of sustained immune-mediated control of tumor growth

Metabolic reprogramming appears to represent a prerequisite for optimal CD8^+^ T cell antitumor activity within the TME (25,26). CD8^+^ T cells from B16F10 tumors were isolated from STZ induced T1D (^STZ^CD8_TIL_) and control (^Cont^CD8_TIL_) mice using MACS (**Figure 4A**) and their purity confirmed by flow cytometry (**Figure 4B**). Subsequent analysis of T cell metabolomics revealed that ^STZ^CD8_TIL_ cells showed a significantly upregulated extracellular acidification rate (ECAR) and oxygen consumption rate (OCR) when compared to ^Cont^CD8_TIL_ cells (**Figure 4C and 4D**). As glucose metabolism is a preferred characteristic for anti-tumor CD8^+^ T cells with regards to their therapeutic efficacy (27–29), we next studied intrinsic T cell expression of transporters and enzymes involved in glucose metabolism (**Figure 4E**). ^STZ^CD8^+^ TILs showed elevated glucose transporter 1 (*glut 1*), monocarboxylate transporter 1 (*mct 1*), and pyruvate kinase M2 (*pkm 2*) expression compared to their respective untreated control CD8^+^ TILs (**Figure 4F**). However, the expression of glucose 6-phosphate dehydrogenase (*g-6-pd*) and lactate dehydrogenase (*ldh*) differed only modestly in ^STZ^CD8^+^ TILs vs. control CD8^+^ TILs. Insulin treatment of STZ induced tumor bearing mice resulted in CD8^+^ TILs with reduced expression of *glut 1*, *mct 1* and *pkm 2* expression vs. ^STZ^CD8^+^ TILs in all four tumor models evaluated in this study. Expression of pyruvate carboxylase 1 (*pcx 1*), which is involved in the conversion of pyruvate (an end-product of glycolysis) into oxaloacetate, was significantly elevated in ^STZ^CD8^+^ TILs isolated from B16F10 and LLC, but not 4T1 and CT26 tumors. Insulin treatment of STZ induced tumor-bearing mice resulted in reduced expression of *pcx 1* in CD8^+^ TILs (vs. ^STZ^CD8^+^ TILs) isolated from B16F10 and LLC, but not 4T1 and CT26 tumors. Isocitrate dehydrogenase 2 (*idh 2*) expression was found differentially enhanced in ^STZ^CD8^+^ TILs from B16F10, 4T1 and LLC (but not CT26) vs. comparable TIL isolated from control or insulin + STZ-treated tumor bearing hosts. These results support divergent expression patterns for *pcx 1* and *idh 2* gene expression in ^STZ^CD8^+^ TIL across the various tumor models. In contrast, transcript levels for enzymes involved in the TCA cycle, such as pyruvate dehydrogenase kinase 1 (*pdk 1*), pyruvate dehydrogenase kinase 3 (*pdk 3*), isocitrate dehydrogenase 1 (*idh 1*) and fumarate hydratase 1 (*fh 1*) were found comparable in CD8^+^ TILs isolated from tumors harvested from untreated (control), STZ treated or STZ+insulin treated mice.

**Figure 4.**
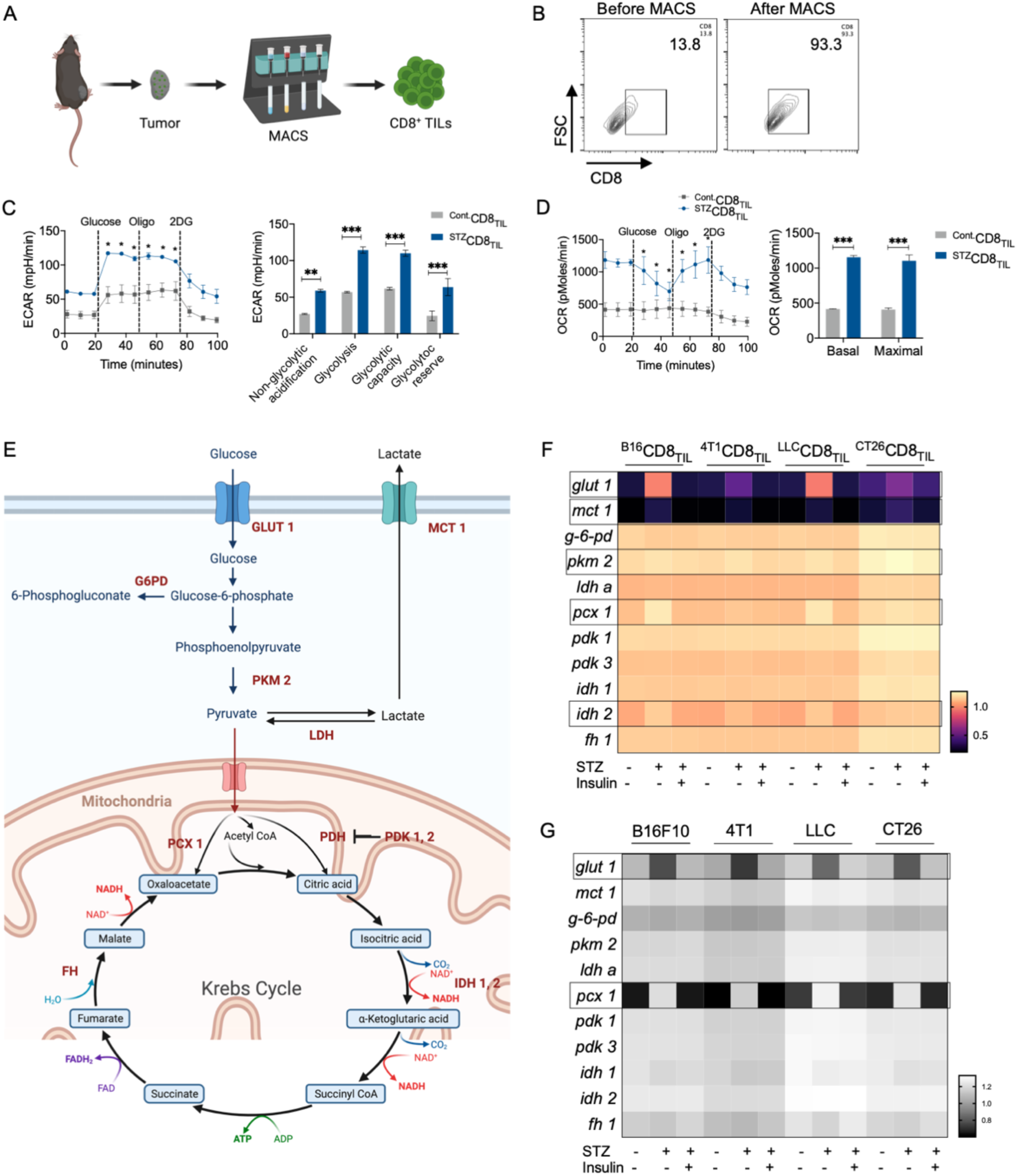
T1D metabolically reprograms CD8^+^ T lymphocytes in support of sustained immune-mediated control of tumor growth. (**A)** Schematic overview for magnetic-activated cell sorting (MACS) of CD8^+^ TIL. (**B)** Flow cytometry contour plots of CD8^+^ T cells before and after MACS sorting. (**C)** Extracellular acidification rate (ECAR) and (**D)** Oxygen consumption rate (OCR) of CD8^+^ TILs isolated from B16F10 tumors (n=6 mice/group) from untreated and STZ induced T1D mice. (**E)** Schematic overview of glycolysis and Krebs cycle along with the pathway associated enzymes analyzed (in red). Gene expression profiles of *glut 1* and *mct 1* transporters, *g-6-pd* and *pkm 2*, enzymes involved in glycolysis, *ldh a*, involved in pyruvate to lactate conversion, *pcx 1*, *idh 1*, *idh 2*, *fh 1*, involved in Krebs cycle, *pdk 1* and *pdk 3,* inhibitor of pyruvate dehydrogenase (PDH), with *β actin* as a control gene product, were measured by RT-PCR in (**F)** CD8^+^ TILs (n=6) or (**G)** tumor cells. Two-way ANOVA was performed to analyze the significance or intergroup differences. (*P ≤ 0.05, **P ≤ 0.01, ***P ≤ 0.001, ****P ≤ 0.0001)

In contrast, B16F10, 4T1, LLC and CT26 tumor cells isolated from STZ induced T1D mice displayed decreased expression of *glut 1* and *pcx 1*, when compared to their respective counterparts in the control and STZ+insulin treated groups (**Figure 4G**). No significant alterations in expression of *mct 1, g-6-pd, pkm 2, ldh a, pdk 1, pdk 3, idh 1, idh 2 and fh1* were noted in tumor cells based on the host treatment regimen. These findings suggest that pre-existent T1D leads to selective glycolytic benefits in CD8^+^ TIL vs. tumor cells, with this metabolic advantage in ^STZ^CD8_TIL_ cells yielding sustained antitumor activity in association with superior tumor growth control in vivo.

### 5. Pre-existent T1D in the tumor bearing host is associated with enrichment in CD8^+^IGF1^+^IGFR^+^ T cells in blood, TDLN and tumor

Given the importance of T1D-associated metabolic reprogramming of CD8^+^ T cells in restricting tumor growth, we next aimed to decipher the underlying signaling mechanisms associated with improved immune-mediated control of tumor growth. Insulin-like growth factor 1 (*igf1*) and insulin-like growth factor 1 receptor (*igf1R*) expression was significantly elevated in ^STZ^CD8^+^ TILs from B16F10, 4T1, LLC and CT26 tumors, with B16F10 TILs exhibiting the most profound degree of overexpression (**Figure 5A**). However, both C57BL/6J and BALB/c mice, after STZ treatment and under tumor-free conditions exhibited significantly reduced serum IGF1 concentrations, which could be normalized following insulin treatment **(Figure S1C)**. Insulin like growth factor 2 receptor (*igf2R*) showed no significant alteration among the different groups of CD8^+^ TILs across all tumor types. Insulin receptor (*insulinR*) was found to be downregulated in TILs from B16F10 melanomas (but not other tumors in our study) established in STZ induced T1D mice. Conversely, tumor cells recovered from STZ induced T1D mice displayed diminished *igf1*, *igf1R* and *insulinR* expression when compared to respective untreated (control) or STZ+insulin treated tumor-bearing cohorts (**Figure 5B**). In contrast, *igf2R* expression remained unchanged, and *igf1 and igf1R* expression was significantly upregulated, in ^STZ^CD8_TDLN_ vs. ^Cont.^CD8_TDLN_ isolated from B16F10 tumors(**Figure 5C**). These metabolic T1D-associated alterations in TDLN T cells were reversed by treatment with insulin and were not observed in (non-tumor) LNs.

**Figure 5.**
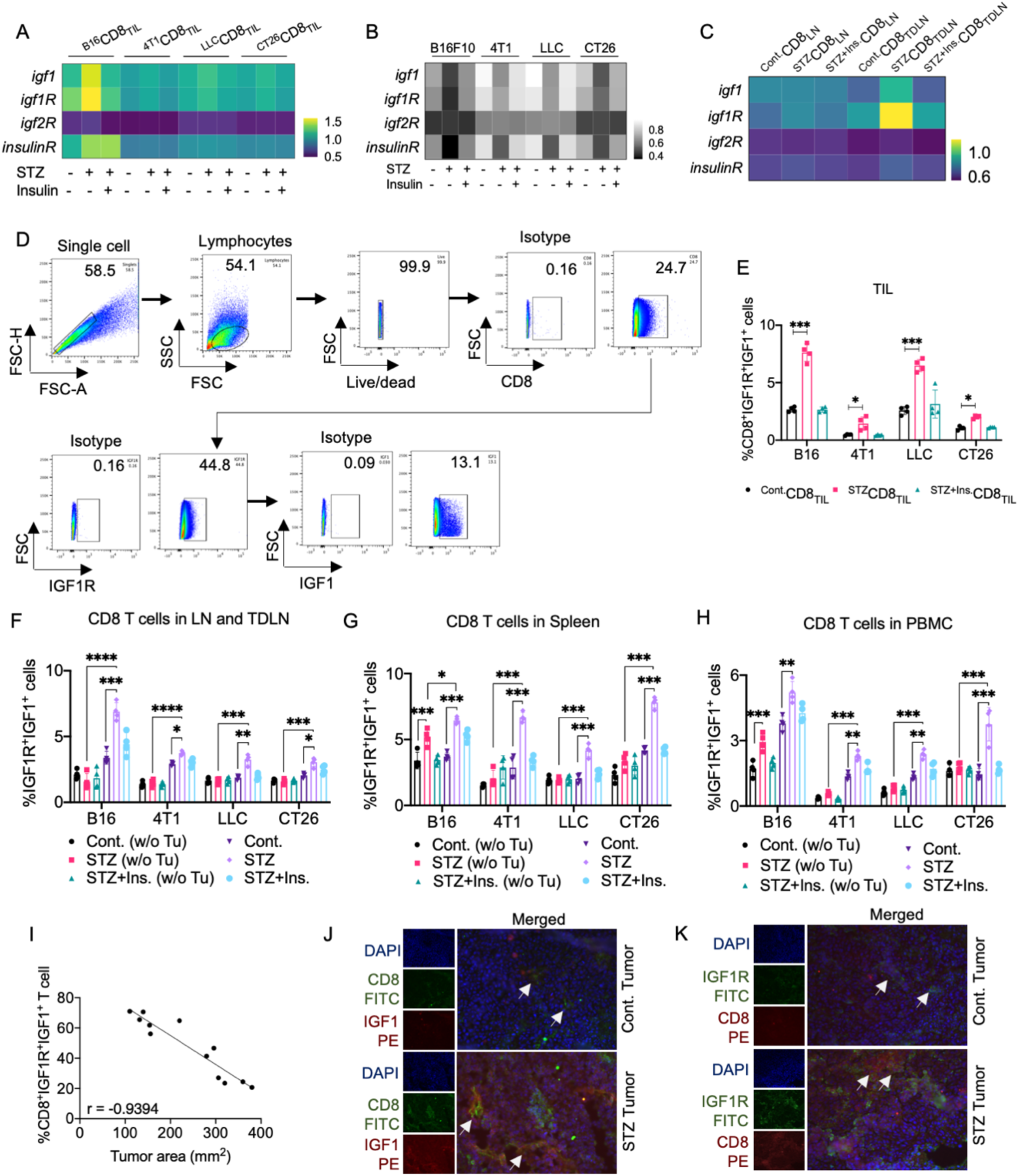
Pre-existent T1D in the tumor bearing host is associated with enrichment in CD8^+^IGF1^+^IGFR^+^ T cells in blood, TDLN and tumor. Expression of *igf1, igf1R, igf2R* and *insulinR* was analyzed by RT-PCR from (**A**) MACS sorted CD8^+^ TILs, (**B**) tumor cells and (**C**) MACS sorted CD8^+^ T cells isolated from TDLNs (n=6 mice/group). (**D)** Flow cytometry gating and **(E)** percentage of CD8^+^IGF1^+^IGF1R^+^ cells in tumors (n=4). CD8^+^IGF1^+^IGF1R^+^ cells in **(F)** LNs and TDLNs, **(G)** spleen and **(H)** PBMCs (n=4 mice/group). **(I)** Correlation analysis between tumor area and the frequency of CD8^+^IGF1^+^IGF1R^+^ T cells in tumors. **(J)** Colocalization of CD8 and IGF1 (arrow marked) in B16F10 tumor sections harvested from control vs. STZ induced T1D mice. **(K)** Colocalization of CD8 and IGF1R (arrow marked) in B16F10 tumor sections harvested from control vs. STZ induced T1D mice. Two-way ANOVA was performed to test for significance across groups. (*P ≤ 0.05, **P ≤ 0.01, ***P ≤ 0.001, ****P ≤ 0.0001)

Co-expression of IGF1R and IGF1 on CD8^+^ TILs was then analyzed by flow cytometry in the various tumor models (**Figure 5D**). ^STZ^CD8_TIL_ were enriched in the CD8^+^IGF1R^+^IGF1^+^ T cell subpopulation when compared to ^Cont.^CD8_TIL_ in all 4 tumor models (**Figure 5E**). Insulin treatment of STZ induced T1D tumor bearing mice yielded ^STZ+Ins.^CD8_TIL_ with significantly reduced frequencies of CD8^+^IGF1R^+^IGF1^+^ TILs vs. ^Cont.^CD8_TIL_ and ^STZ^CD8_TIL_ across all models Exploration of CD8^+^IGF1R^+^IGF1^+^ cell population was further extended to LNs, TDLNs, spleens and PBMCs. LNs isolated from tumor-free untreated, STZ treated and STZ+insulin treated mice did not exhibit significant difference in CD8^+^IGF1R^+^IGF1^+^ T cell populations (**Figure 5F** **and S5**). However, the CD8^+^IGF1R^+^IGF1^+^ T cell subpopulation was elevated in TDLNs of STZ treated tumor bearing mice, regardless of the tumor model studied. Spleens of STZ (but not STZ+insulin) treated tumor bearing mice also showed upregulated frequencies of CD8^+^IGF1R^+^IGF1^+^ T cells when compared to spleens harvested from control groups (**Figure 5G** **and S6)**. Similar results were obtained when analyzing CD8^+^IGF1R^+^IGF1^+^ T cell population in PBMC in all four tumor models (**Figure 5H** **and S7)**. Notably, CD8^+^IGF1R^+^IGF1^+^ cells were only increased in tumor bearing mice, with the percentage of CD8^+^IGF1^+^IGF1R^+^ T cells amongst total TILs determined to be inversely correlated with tumor size (**Figure 5I**). Moreover, IFM analyses of B16F10 tumor sections from STZ treated T1D mice showed increased colocalization of IGF1 and IGF1R on CD8^+^ TIL when compared to untreated B16F10 tumor sections (**Figure 5J and 5K**).

### 6. IGF1-IGF1R-mTOR signaling influences T1D associated metabolic activation of CD8^+^ T cells in the tumor bearing host

As CD8^+^ T cells in T1D tumor bearing mice exhibit selective coordinate upregulation of intrinsic IGF1 and IGF1R expression,, next we studied the associated downstream signaling pathway in these T effector cells. ^STZ^CD8_TIL_ cells isolated from B16F10 melanomas expressed significantly elevated levels of phospho-AKT and phospho-mTORC1 when compared to ^Cont^CD8_TIL_ cells (**Figure 6A**), with these changes selectively occurring in CD8^+^ TIL but not tumor cells **(Figure S8)**. *In silico* protein-protein interactome analyses with STRING revealed a operational connection between IGF1, IGF1R, AKT and mTOR (**Figure 6B**). To confirm the involvement of mTORC1 with upregulated IGF1/IGF1R activity in the activation of CD8^+^ T cells, *in vitro* experiments were performed with MACS CD8^+^ T cells isolated from the TDLNs of STZ induced T1D mice bearing B16 F10 melanomas vs. control tumor bearing mice (**Figure 6C**). Recombinant IGF1 (rIGF1) was used as an IGF1R activator, with small interfering RNA against IGF1R (siIGF1R) used to block IGF1R expression. Rapamycin was used to block mTORC1-mediated signaling. ^Cont.^CD8^+^ T cells and ^STZ^CD8^+^ T cells from TDLNs were treated with either rIGF1 or siIGF1R or rapamycin alone or in combinations. We observed that ^STZ^CD8^+^ T cells exhibited significant increases in expression of *igf1R, glut1, gzmB and ifnγ* after treatment with rIGF1 (**Figure 6D**). However, the expression of these genes was significantly downregulated when ^STZ^CD8^+^ T cells were treated with either siIGF1R or rapamycin, or a combination of both agents. Flow cytometry analysis confirmed that rIGF1 treatment of ^STZ^CD8^+^ T cells, but not ^Cont.^CD8^+^ T cells, coordinately resulted in significantly upregulated expression of CD69 and IFNγ (**Figure 6E and 6F**). Treatment with either siIGF1R or rapamycin or both agents led to a significant diminishment in CD69 and IFNγ expression by ^STZ^CD8^+^ T cells but not by ^Cont.^CD8^+^ T cells. These results suggest that ^STZ^CD8^+^ T cells from TDLN are more susceptible to targeted antagonism of IGF1R and mTOR than their non-diabetic counterparts.

**Figure 6.**
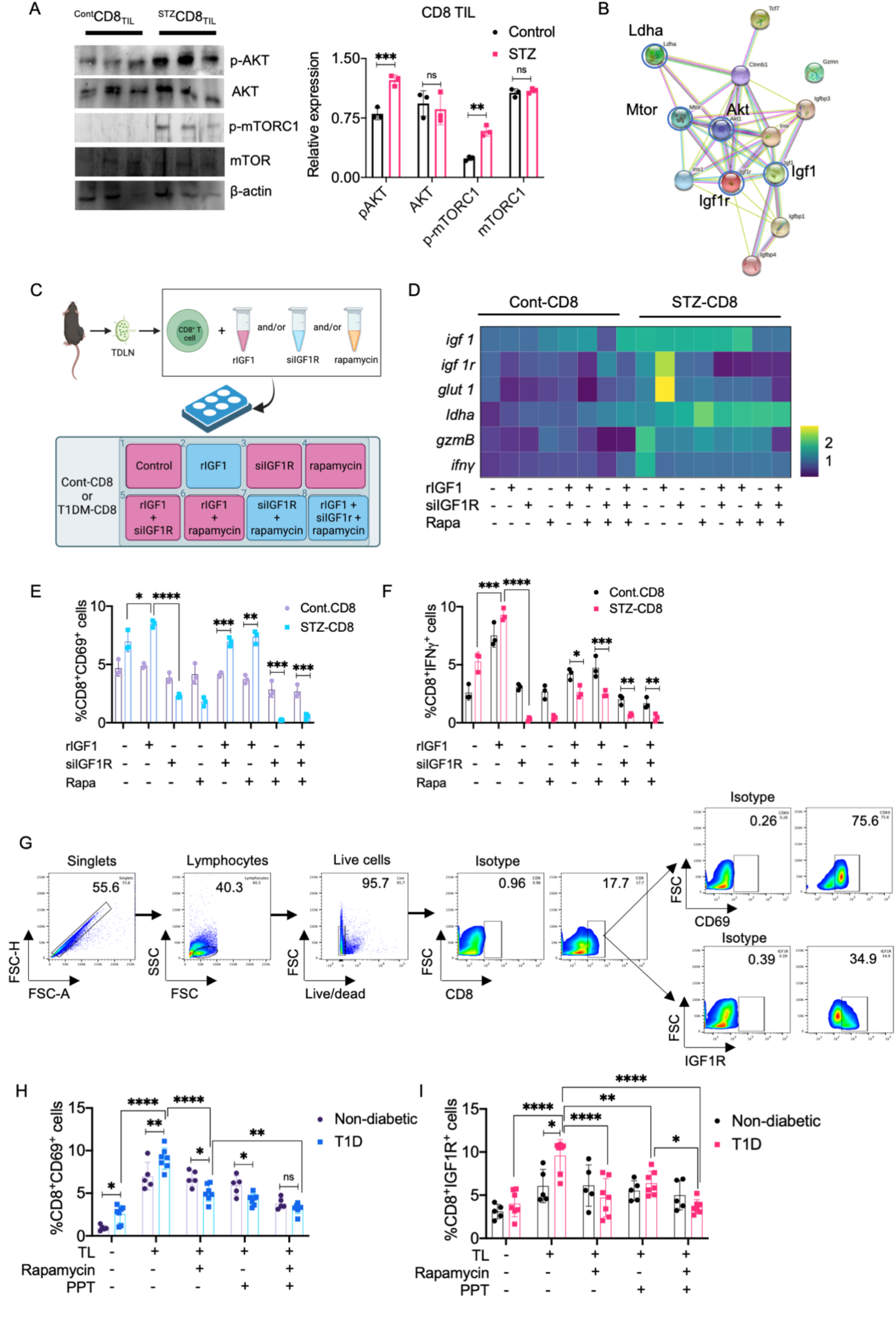
IGF1-IGF1R-mTOR signaling influences T1D associated metabolic activation of CD8^+^ T cells in the tumor bearing host. (**A)** Western blot analysis of pAKT, AKT, mTOR and phosphor-mTORC1 (p-mTORC1) in MACS sorted CD8^+^ TILs (n=3). (**B)** Search tool for the retrieval of interacting genes/proteins (STRING) analysis was used to study signaling interactions between IGF1, IGF1R, InsulinR and mTOR. (**C)** Schematic overview of *in vitro* experiments (n=3) to study cellular signaling in MACS sorted CD8^+^ T cells isolated from TDLN. rIGF1 was used to stimulate the IGF1-IGF1R-mTOR signaling axis, siIGF1R and rapamycin were used to block IGF1R expression and mTORC1, respectively. (**D)** Expression of *igf1, igf1R, glut1, ldha, gzmB* and *ifnγ* following *in vitro* treatment of rIGF1, siIGF1R and rapamycin, either alone or in combination. **(E)** Percent CD8^+^CD69^+^ cells in ^Cont.^CD8^+^ T cells vs. ^STZ^CD8^+^ T cells (n=3). **(F)** Percent CD8^+^IFNγ^+^ cells amongst ^Cont.^CD8^+^ vs. ^STZ^CD8 groups (n=3). **(G)** Flow cytometry gating on CD8^+^, CD69^+^ and IGF1R^+^ cells in T1D patient PBMC. **(H)** Percent CD8^+^CD69^+^ and **(I)** percent CD8^+^CD69^+^ cell population in T1D patient and non-diabetic PBMC samples after in vitro treatment with tumor lysate (TL), rapamycin and picropodophyllotoxin (PPT) alone or in combination. Two-way ANOVA followed by Tukey’s multiple comparison test was performed to test significance of intergroup differences. (*P ≤ 0.05, **P ≤ 0.01, ***P ≤ 0.001, ****P ≤ 0.0001)

To confirm this finding in the clinical setting, T1D patient PBMCs, along with age and sex matched non-diabetic PBMCs, were stimulated in vitro with a lysate derived from the human breast cancer cell line MCF 7 in the absence or presence of rapamycin (mTOR inhibitor) or picropodophyllotoxin (PPT; an IGF1R inhibitor) or both. Flow cytometry was performed to assess CD8^+^CD69^+^ cells and CD8^+^IGF1R^+^ status amongst “responder” T cells (**Figure 6G**). Although the CD8^+^CD69^+^ T cell population was determined to be elevated in the stimulated PBMCs from both T1D and non-T1D responders following stimulation with tumor antigens (**Figure 6H**), such upregulation was most prominent in T1D PBMCs Interestingly, rapamycin or PPT significantly downregulated the frequency of CD8^+^CD69^+^ cells in T1D-PBMCs compared to non-T1D PBMCs, with CD8^+^IGF1R^+^ cells found to be elevated in cultures developed from both T1D and non-T1D (with T1D > non-TID) PBMCs following antigenic stimulation (**Figure 6I**). Co-treatment with rapamycin or PPT or both resulted in a significant reduction in the CD8^+^IGF1R^+^ T cell population from T1D donors vs. treatment with lysate only. These findings support the involvement of an IGF1-IGF1R-mTORC1 axis in the superior activation of (anti-tumor) CD8^+^ T cells in T1D hosts that underlies enhanced tumor growth control *in vivo*.

## Discussion

The present study reports the novel finding that T1D predisposes the host to improved CD8^+^ T cell-mediated control of tumor growth in a generalized, pan-cancer manner (across a range of murine melanoma and carcinoma models). Streptozotocin (STZ) selectively destroys pancreatic beta cells and is widely used to develop models reflective of T1D (18,19). STZ enters cells through the glucose transporter 2 (Glut 2), which is expressed at high levels on pancreatic β cells, where it operationally functions as a glucose sensor(30). STZ induced T1D animals show identical pathological and molecular signatures found in T1D patients (31,32). Intriguingly, a previous *in vivo* study suggested that alloxan-induced T1D in mammary carcinoma bearing rats could lead to the regression of tumors in a majority of the cases (17). In that study, mammary carcinomas were experimentally induced in rats, with animals then treated the with T1D inducing drug alloxan, providing conditions for direct effect of alloxan on cancer cells. We eliminated the possibility for direct anti-tumor effects in our model by discontinuing STZ treatment for 4 weeks prior to tumor inoculation(33). Before tumor cell inoculation, STZ treated mice displayed significantly elevated blood glucose levels and diminished serum insulin levels when compared to untreated and STZ+insulin treated mice. Upon insulin treatment, blood glucose in STZ+insulin animals was normalized, with insulin levels left unchanged (indicative of the permanent destruction of pancreatic beta cells). Importantly, we noted that reduced levels of serum IGF1 in STZ only treated mice were restored to normal levels in insulin+STZ treated mice. IGF1 in circulation primarily derives from hepatocytes upon insulin stimulation. Analyses of other serum cytokines revealed STZ treatment upregulated IL-6 in circulation and that treatment with insulin normalized levels of this chronic inflammatory biomarker. Levels of serum IL-17, IFNγ, TNFα and IL-10 appeared unaltered under STZ +/- insulin treatment conditions.

Significant tumor growth control and extended host survival were observed in STZ induced T1D mice bearing any of 4 unrelated tumors that vary across histologies which was antagonized by treatment with insulin.. In an attempt to evaluate the possible involvement of adaptive immune system in T1D hosts, we extended our study in athymic nude mice lacking a functional adaptive immune system. STZ induced T1D nude mice failed to control melanoma growth or extend survival when compared to untreated melanoma bearing nude mice. STZ induced T1D tumors (B16F10, 4T1, LLC, CT26) were characterized by increased infiltration by CD8^+^ T cells enriched in a polyfunctional proinflammatory phenotype (i.e., expressing elevated levels of CD69, CXCR3, GZMB and IFNγ) and ^STZ^CD8_TIL_ displayed superior anti-tumor cytotoxic activity when compared to ^Ctrl^CD8_TIL_. Enrichment of CD8^+^CD103^+^CD39^+^ cells amongst TIL, the CD8^+^CD69^+^GZMB^+^ subpopulation amongst TDLNs in STZ induced T1D mice bearing B16F10, 4T1, LLC or CT26 tumors is indicative of improved levels of tumor antigen specific CD8^+^ T cells in these diabetic animals. The crucial role of CD8^+^ T cells in tumor growth restriction in T1D mice was confirmed in adoptive CD8^+^ T cell transfer models applied to athymic T1D tumor-bearing recipient mice and via the use of in vivo CD8 T cell depletion in STZ induced T1D tumor models. ^53,54^The superior anti-tumor efficacy of ^STZ^CD8^+^ T cells in our diabetic models may be causally linked to 1.) the co-ordinately low levels of checkpoint molecules (PD1, LAG3) and higher expression of TCF1 associated with a less exhausted TIL effector cell populations and 2.) reduced levels of immunoregulatory cell populations (including regulatory T cells and MDSCs) found in the TME of T1D host animals.

Our study findings suggest that pre-existing T1D, in the absence of insulin treatment or diet alterations, might have a protective role in cancer. It might be noted that insulin has an important role on cancer cell metabolism where it has been found to be mitogenic, but not carcinogenic (34,35). Studies suggest that during and after the differentiation of naive CD8^+^ T cells into effector T cells following antigen cross-presentation from antigen-presenting cells, CD8^+^ T cells rely on aerobic glycolysis, similar to the Warburg effect observed in cancer cells (27,36). We found that ^STZ^CD8_TIL_ cells from B16F10 tumor bearing mice exhibit significantly elevated glycolysis and Krebs cycle activation when compared to ^Cont.^CD8_TIL_ cells. Higher expression of *glut 1, mct 1, pkm 2*, *pcx 1 and idh 2* in ^STZ^CD8_TIL_ in tumor bearing hosts further suggests intrinsic metabolic activation of CD8^+^ TILs under diabetogenic conditions. Whereas, insulin treatment of STZ induced T1D negatively impacts metabolic genes in CD8^+^ TILs recovered from all four tumor models examined in this study. Tumor cells in STZ induced mice on the other hand, showed diminished *glut 1* expression and increased *pcx 1* expression indicative of repressed glycolytic activity and slowed proliferative capacity. Our observations suggest that within the context of the T1D TME, CD8^+^ TILs gain a metabolic advantage over tumor cells in association with enhanced anti-tumor CD8^+^ T cell infiltration, activation and sustained effector function resulting in improved tumor growth control.

Further studies revealed that CD8^+^ TILs, but not tumor cells, expressed significantly higher IGF1 and IGF1R mRNA and protein levels. The frequency of CD8^+^IGF1^+^IGF1R^+^ cells was significantly higher in TILs, TDLNs, spleens and PBMCs of STZ induced T1D tumor-bearing mice but not in non-diabetic tumor hosts. Interestingly, the proportion of CD8^+^IGF1^+^IGF1R^+^ cells amongst TIL was found to be inversely correlated with tumor size. This observation suggests a crucial role for this effector cell population in restricting tumor growth in T1D hosts. Untreated, STZ treated and STZ+insulin treated hosts without tumor, however, showed no significant change in the frequency of the CD8^+^IGF1^+^IGF1R^+^ population in LNs, spleens and PBMCs, with the notable exception of spleen and PBMC in the B16F10 melanoma model.

IGF1 and IGF1R shares significant structural homology with insulin and insulin receptor (IR) respectively. Moreover, insulin and IGF1 are involved in many overlapping signaling pathways (37,38). Although a variety of cells secrete IGF1, the major source of systemic IGF1 is the liver (39). Hepatocytes secrete IGF1 upon stimulation with insulin and growth hormone (GH) (40). T1D patients with very low or undetectable levels of insulin also exhibit reduced plasma IGF1 levels (41). Reduced serum insulin and IGF1 concentrations in STZ induced T1D mice may trigger enhanced expression of IGF1 and IGF1R on CD8^+^ T cells upon cognate (tumor) antigen stimulation. This may trigger autocrine IGF1-IGF1R signaling in CD8^+^ T cells, which may be responsible for the enhanced metabolic activation and cytotoxic potential of these cells. Acute fasting has been reported to reduce systemic IGF1 levels, to boost systemic CD8^+^ T cell metabolism, and to enhance CTL-mediated cytotoxicity (42). Glucose transporter 2 (Glut-2) which is predominantly expressed by pancreatic β cells, can sense a rise in blood glucose, which in turn stimulates insulin secretion (43). During fasting, blood glucose levels are normalized there is little to no insulin in circulation. These conditions may result in decreased systemic IGF1 levels, which in turn could force CD8^+^ T cells to activate autocrine IGF1 production and *cis/trans* activation of IGF1R-mediated signaling. *In vitro* studies have previously suggested that activated CD8^+^ T cells produced significant quantities of IGF1 (44). These cumulative findings support the hypothesis that autocrine IGF1 signaling may be activated in CD8^+^ T cells when systemic IGF1 is downregulated, as in the case of tumor-bearing T1D hosts. IGF1 is a potent mitogen, and high systemic IGF1 levels are associated with poor prognosis in several types of cancers (45). IGF1-IGF1R signaling in tumor cells results in enhanced cell proliferation, metabolic activation, escape from programmed cell death, and metastasis. However, clinical trials using IGF1R targeting antagonists or monoclonal antibodies have shown only limited success to date (46). IGF1 can also induce CD8^+^ T cell activation, proliferation, and chemotaxis through the IGF1R(47). Notably, high IGF1R expression is required for leukemia-initiating cell activity in T-cell acute lymphoblastic leukemia (T-ALL) (48). IGF1R is also critical for regulating Th17-Treg balance in autoimmune diseases (49). ^73^Considering the differential and context-dependent effects of IGF1-IGF1R signaling across various cells and diseases, targeted modulation of such circuitry in effector CD8^+^ T cells might be exploited for therapeutic gain in the setting of adoptive T cell-based therapy of cancer.

In B16F10 melanoma models, we observed a significant upregulation of pAKT and p-mTORC1 in ^STZ^CD8_TIL_ cells vs. ^Cont.^CD8_TIL_ cells. Protein-protein interaction network studies further revealed an interconnection between IGF1, IGF1R, AKT and mTOR. We hypothesised that metabolic upregulation and enhanced effector function in CD8^+^ T cells from STZ induced T1D mice might be mediated through the IGF1R-mTOR signaling axis. To test our hypothesis, we isolated CD8^+^ T cells from TDLNs of STZ induced T1D vs. control B16F10 melanoma bearing mice. Isolated ^Cont^.CD8 T cells and ^STZ^CD8 T cells were treated with either recombinant IGF1 (rIGF1) or with silencing RNA against IGF1R (siIGF1R) or with rapamycin (mTOR inhibitor) alone or in combination. rIGF1 significantly enhanced activation markers on ^STZ^CD8 T cells but not ^Cont^CD8 T cells. ^STZ^CD8 T cells were determined to be more sensitive to treatment with siIGF1R or rapamycin or the combination of these agents. Our results clearly indicate that the IGF1R-mTOR signaling axis underlies the enhanced effector function of CD8^+^ T cells in STZ induced T1D tumor bearing mice. Furthermore, we validated this core paradigm in clinical samples isolated from T1D patients. In these in vitro analyses. we stimulated PBMCs from T1D vs. non-diabetic patients with a source of tumor antigen (i.e., tumor lysate). We observed significant upregulation in responder populations of CD8^+^CD69^+^ T cells and CD8^+^IGF1R^+^ T cells in cultures developed from T1D PBMCs vs. non-diabetic PBMCs. Targeted antagonism of IGF1R or mTOR or both resulted in a significant reduction in these two CD8^+^ T cells subpopulations in T1D PBMC cultures. In vitro stimulation with tumor antigens stimulated greater CD8^+^ T cell activation from T1D vs. non-diabetic patients. CD8^+^ T cells from T1D patients were more sensitive to the inhibitory effects of IGF1R blockers or mTOR antagonists. This observation strengthens our hypothesis that the IGF1R-mTOR signaling axis in CD8^+^ T cells in T1D hosts is critical to tumor growth control. Sustained CD8^+^ T cell fate/function are essential for the success of cancer immunotherapy, with the success of anti-PD1 therapy link to the ability of this agent to prevent the exhaustion and premature death of ant-tumor CD8^+^ T cells (50). Very recently, *cis* B7:CD28 costimulatory interactions in CD8^+^ T cells have been demonstrated to promote anti-tumor immunity, implying that CD8^+^ T cells could boost themselves in an autocrine manner (51). In a similar manner, intrinsic IGF1R-mTOR signaling in CD8^+^ T cells may be important to the superior anti-tumor action of these effector cells in T1D patients. Interestingly, a fraction of cancer patients undergoing immune checkpoint blockade therapy (ICBT) develop immune-related adverse effects (irAEs), among which T1D is very common (52). It has not yet been documented whether cancer patients who develop T1D during ICBT have a better clinical prognosis. We would hypothesize that development of T1D may enable IGF1/IGF1R/mTOR-dependent anti-tumor immunity post-ICBT therapy. ^77^The IGF1R-mTOR signaling axis could also be exploited within the context of CAR-T cell-based cancer immunotherapy, where T cell anergy and exhaustion remain major challenges to the therapeutic efficacy of this approach in the solid cancer setting (53). Future work will define optimal targeting agents to agonize IGF1R-mTOR signaling in anti-tumor CD8^+^ T cells for sustained therapeutic benefit in cancer patients.

## Limitations of the study

In this study we used streptozotocin induced T1D mouse model. Though programming in the NOD mouse model mimics human T1D initiation and progression, the lack of standardised syngeneic tumor models in NOD mice led to our choice to use low dose streptozotocin to induce T1D in C57BL/6 and BALB/c mice for which a broad range of transplantable models were available for *in vivo* modeling. Establishing syngeneic tumor models in NOD mice will allow for future validation testing of the key postulates and findings reported in this manuscript. Other types of cancer models beyond translantable systems, including (spontaneous/conditional tumor) GEMM should also be evaluated in future to test the generality of our findings. For the human PBMC studies outlined in this report, the T1D patient sample size was small due to limited availability of these donors in our community. Further validation with larger cohorts of donors is warranted.

## Ethical approval

Experiments using C57BL/6J and BALB/c mice were approved by the Institutional Animal Ethics Committee of the Chittaranjan National Cancer Institute, Kolkata, India (Approval no. IAEC-1774/RB-17/2017/4), registered under CPCSEA, India. Experiments using nude mice were approved by the Institutional Animal Ethics Committee, Central Drug Research Institute, Lucknow, India (Approval No. IAEC/2019/78/Renew-0/Dated-05/04/2019). Experiments using type 1 diabetic patients blood samples were approved by the Institutional Ethics Committee, DDRF, Kolkata (Approval No. DDRF-IEC-74517-RB-2022-1), registered under the Central Drug Standard Control Organization (CDSCO), India.

## Acknowledgements

We thank Director, Chittaranjan National Cancer Institute (CNCI), Kolkata, India, Director, CSIR-Indian Institute of Chemical Biology (CSIR-IICB), Kolkata, India, and CSIR-Central Drug Research Institute (CSIR-CDRI), Lucknow, India for providing instrumental facilities in their respective institutes. We also thank Dr. Abhijit Rakshit, Head, Animal Facility, CNCI, Kolkata, India. We would like to thank Mendeley Ltd., Elsevier for providing a free software tool for arranging references.

## Authors’ contributions

Conceptualization, A.B. and A.S.; Methodology, A.S., A.B., A.V. and P.P.; Validation, A.S., A. Bhuniya., I.G., S.D., S.B., A.Saha.; Formal analysis, A.S. and A.B.; Investigation, A.S., S.D., S.B., M.C., A.V., P.P., J.S. and J.D.; Resources, S.S.R., S.R, D.B., S.B., R.B. and D.D.; Writing-original draft, A.S.; Writing-Review & Editing, A.B., R.B. and W.J.S.; Visualization, A.S.; Supervision, A.B. and R.B; Project administration, R.B; Funding Acquisition, A.S., A.B., S.B., D.D. and R.B.

## Materials and Methods

### Patients and samples

Random blood from type-1 diabetic (T1D) patients (n=7) with no other disease reported in last one month, were obtained from Doyen Diagnostics and Research Foundation (DDRF), Kolkata, India, following obtaining their written consent. Ethical approval of this study was obtained from Institutional Ethics Committee, DDRF, Kolkata (Approval No. DDRF-IEC-74517-RB-2022-1), registered under the Central Drug Standard Control Organization (CDSCO), India. Blood samples from age-and sex-matched healthy volunteers (n=5) with no reported disease for past one month were collected as control after obtaining their written consent. Blood glucose, HbA1c, glutamic acid decarboxylase (GAD65) antibody and anti-insulin antibody was measured in plasma. T1D patient and healthy volunteer data was summarised in **Supplementary Table ST1 and ST2**. All the study were strictly adhered to the Declaration of Helsinki. Patients and healthy volunteers included in this study are of South Asian ancestry.

### Mice

Male C57BL/6J mice, male BALB/c mice and female BALB/c (age, 4–6 weeks, body weight, 25 g on average) were obtained from and maintained at the Institutional Animal Facility, Chittaranjan National Cancer Institute (CNCI), Kolkata. Nude Crl:CD1-Foxn1^nu^ mice (age, 4–6 weeks, body weight, 25 g on average) were obtained from and maintained at the Animal Facility, CSIR-Central Drug Research Institute (CSIR-CDRI), Lucknow. Autoclaved dry pellet diet and water were provided *ad libitum.* Animals were maintained and treated according to the guidelines established by the CPCSEA, Govt. of India, following approval from the Institutional Animal Ethics Committee (IAEC), CNCI, Kolkata (Approval no. IAEC-1774/RB-17/2017/4), and IAEC, CDRI, Lucknow (Approval No. IAEC/2019/78/Renew-0/Dated-05/04/2019) respectively.

### Cell lines and reagents

B16F10 (melanoma), 4T1 (breast carcinoma), LLC (lung carcinoma), CT26 (colon carcinoma) and S180 (sarcoma) cell lines were obtained from the National Center for Cell Sciences (Pune, India). B16F10 and LLC cells were cultured in high-glucose DMEM (Himedia) supplemented with 10% (v/v) heat-inactivated FBS (Himedia), 2 mM L-glutamine, penicillin (1000 U/mL), and streptomycin (10 mg/mL) (Himedia). 4T1 and CT26 and S180 cells were cultured in RPMI-1640 (Himedia) supplemented with 10% (v/v) heat-inactivated FBS (Himedia), 2 mM L-glutamine, penicillin (1000 U/mL), and streptomycin (10 mg/mL) (Himedia). Cells were incubated in a CO_2_ incubator (Thermo Fisher Scientific) at 37 °C with 5% CO_2_ and 95% humidity. A list of reagents are mentioned in **Supplementary Table ST3**.

### Murine tumor model for type 1 diabetes

T1D was induced in 4-6 hrs fasted mice by intraperitoneal (i.p.) injection of low dose streptozotocin (Millipore) at 50 mg/kg for five consecutive days, dissolved in monohydrate Na-citrate buffer pH 4.5 (54) under anaesthesia as per the CPCSEA guidelines. Anesthesia was induced using 3.5% isoflurane for four minutes in medical air enriched with oxygen (air:oxygen 3:1). The control group received vehicle. Sucrose water (10%) was given overnight to avoid sudden hypoglycemia post injection. Mice were kept for four weeks with an autoclaved dry pellet diet and water *ad libitum* for proper diabetes induction. After four weeks, only those mice with blood glucose between 250-350 mg/dl were taken for next stage of experiment. Non-diabetic control group had a mean blood glucose of 100 mg/dl. One group of T1D induced mice received 1UI/kg insulin i.p. twice daily, from day 1, throughout the experiment. Blood sugar was measured, after four weeks, using the glucose oxidase-peroxidase (GOD-POD) method following the manufacturer’s protocol (Autospan Glucose measuring kit, India) to evaluate hyperglycemia.

### Solid tumor development

Solid tumors were developed in mice after four weeks of T1D induction, by inoculation of tumor (5×10^5^) cells s.c. on the right flank of non-diabetic, STZ and STZ+insulin group of mice. B16F10 and LLC tumors were induced in male C57BL/6J mice. CT-26 tumors were induced in male BALB/c mice. 4T1 tumors were induced in female BALB/c mice in mammary fat pad. B16F10 tumors were also induced in male Crl:CD1-Foxn1^nu^ nude mice. After palpable tumor formation, tumor areas (mm^2^) were measured regularly up to day 28 for B16F10, 4T1, LLC and CT-26. B16F10 tumors in nude mice were measured up to day 19 following the formula (length × width). Animal health, body weight (g), and blood sugar (mg/dL) of both groups of mice were monitored throughout the experiment. Tumor-bearing mice were euthanized by an overdose of ketamine HCL (160 mg/kg) + xylazine (20 mg/kg) intraperitoneally (i.p.) on days 31 and 19, respectively, as per the CPCSEA guidelines. Tumors, spleens, tumor-draining lymph nodes (TDLN) and blood were collected for downstream analyses.

### Isolation of CD8^+^ T cells from solid tumors and tumor draining lymph nodes

After surgical resection, the tumor samples were rinsed with antibiotic-containing media and minced with scissors under sterile conditions. Enzymatic digestion was performed using 1500 U/ml collagenase IV (Merck) in RPMI, followed by mechanical shaking. Single cells were obtained by passing the enzyme-digested tumor cell suspension through a 70 µm cell strainer (Corning). Tumor-infiltrating lymphocytes (TILs) were purified from buffy coats by centrifugation of digested single-cell suspension over a lymphocyte separation medium made of polysaccharose and diatrizoic acid dihydrate according to the manufacturer’s protocol (Himedia). CD8^+^ T cells from lymph nodes and TILs of B16F10 melanoma-bearing T1D and non-diabetic C57BL/6J mice were isolated by Magnetic-Activated Cell Sorting (MACS) using Anti-Mouse CD3 followed by BD iMag Anti-Mouse CD8a DM (BD Biosciences) following the manufacturer’s protocol. Purity was checked by flow cytometry using anti-mouse CD8a Antibody (BioLegend).

### Adoptive transfer of CD8^+^ T cells to nude mice

CD8^+^ T cells (>90% pure by flow cytometry), MACS sorted from lymph nodes of STZ treated and untreated B16F10 tumor bearing C57BL/6J mice were adoptively transferred (2 × 10^5^ cells) to nude tumor-bearing mice groups through tail vein (i.v.) injection under anesthesia with Ketamine HCl (80 mg/kg) + xylazine (10 mg/kg) i.p. as per the CPCSEA guidelines. CD8^+^ T cells isolated from untreated B16F10 tumor bearing C57BL/6J mice were transferred to untreated nude mice with B16F10 tumor. CD8^+^ T cells from STZ treated B16F10 tumor bearing C57BL/6J mice were transferred to STZ treated nude mice with B16F10 tumor. Tumor progression was monitored in nude mice.

### CD8^+^ T cell depletion and CD8^+^ T cell adoptive transfer

This experiment was conducted to observe how CD8^+^ T cell depletion alters tumor growth in STZ treated mice, also to observe whether adoptive transfer of CD8^+^ T cells from untreated mice to STZ treated mice, and vice versa, cause tumor growth restriction. C57BL/6J mice were divided into eight groups (n=5), four groups received STZ treatment (50 mg/kg for consecutive 5 days) while other four remains untreated. Anti CD8a antibody (100μg/50μl were injected i.p on day -1 and 6 in three untreated and three STZ treated groups. B16F10 tumors were inoculated on day 1 in all groups. On day 7, one untreated and one STZ treated group received adoptive T cell from TDLN of untreated tumor bearing mice. One untreated and one STZ treated group received adoptive T cell from TDLN of STZ treated tumor bearing mice. Tumor area were measured on day 21.

### LDH release assay

Cytotoxicity of MACS-sorted CD8^+^ T cells from B16F10, 4T1, LLC and CT26 tumors were measured by LDH release assay. Sorted cells were cultured *in vitro* against B16F10, 4T1, LLC and CT26 respectively. Cytotoxicity was evaluated by measuring LDH release according to the manufacturer’s protocol (Roche). S180 cell line was used as the unrelated control. The tumor cell: CD8^+^ T cell ratios used in the experiment were 1:10, 1:25, and 1:50. Percent cytotoxicity was calculated using the following formula,

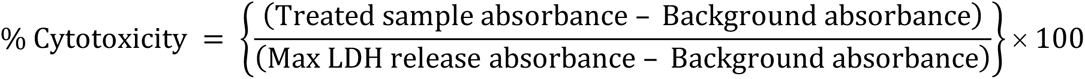

### Extracellular acidification rate (ECAR) and oxygen consumption rate (OCR) quantification

MACS sorted intra-tumoral CD8^+^ T cells were analyzed for glycolysis and Krebs cycle by measuring ECAR and OCR, respectively, in a Seahorse XFe24 Analyzer (Agilent) using a Seahorse XF Glycolytic Rate Assay Kit (Agilent) following the manufacturer’s protocol. Briefly, ECAR and OCR were measured at the basal level, followed by sequential injections of glucose, oligomycin, and 2-DG.

### RNA isolation, cDNA synthesis and RT-PCR

Total cellular RNAs from MACS sorted intra-tumoral CD8^+^ T cells was isolated using TRIzol (Invitrogen). Random hexamers were used to generate the corresponding cDNA using the First Strand cDNA Synthesis Kit (Thermo Scientific) according to the manufacturer’s protocol. Amplification of the target genes was performed using 2x green mix (Promega) with gene-specific primers following the manufacturer’s protocol. RT-PCR products were ran into agarose gel and band intensities were quantified by using ImageJ software. Target gene expressions were normalized relative to the housekeeping gene expression. Statistical analysis was performed to assess the significance of observed differences in target gene expression. Primer sequences and melting temperatures (T_m_) are listed in **Supplementary Table ST4**.

### Western blot

Western blot analysis of mTOR and phospho m-TORC1 from MACS sorted intra-tumoral CD8^+^ T cells and tumor cells was performed. Tumor cells and/or CD8^+^ T cells were lysed in RIPA buffer, and tumor lysates (50 µg) were separated on 10% sodium dodecyl sulfate-polyacrylamide gel using a BioRad apparatus and transferred onto a nitrocellulose membrane (Himedia) for western blotting. Incubation was performed with different primary antibodies after blocking with 5% BSA (Sisco Research Laboratory). After washing, the blots were incubated with horseradish peroxidase-conjugated secondary antibody for 2h at room temperature. The ECL Western Blotting Substrate Kit (Advansta) was used to develop protein bands, which were visualized using Bio-Rad Chemi-Doc XRS^+^.

### Isolation of peripheral blood mononuclear cells (PBMCs)

PBMCs from mice, patient and healthy volunteers’ blood were separated by density gradient centrifugation following the manufacturer’s protocol (Cat# HiSep LSM 1084 and Cat# HiSep LSM 1077, Himedia).

### Flow-cytometry

Flow cytometry acquisitions and analyses were performed using a BD LSRFortessa Cell Analyzer (BD Bioscience) and BD FACSCalibur (BD Bioscience). The data were generated by a fluorometric analysis of 50,000 or 100,000 events per experimental requirement. A list of antibodies used is provided in Key Resources Table. Briefly, for cell surface staining, cells were incubated with anti-mouse monoclonal fluorescently labelled antibodies (specific and isotype-matched controls) for 30 min at 4°C in the dark. 1:500 dilution was used for all antibodies used in flow cytometry. Cells were washed twice with FACS buffer (0.1% BSA in PBS) and fixed in 1% paraformaldehyde before acquisition. Intracellular molecules were stained with anti-mouse fluorescence-labelled monoclonal antibodies using Cytofix/Cytoperm buffer (BD Biosciences), according to the manufacturer’s protocol. Analyses were performed using FlowJo v.10 software (BD Biosciences).

### Enzyme linked immunosorbent assay (ELISA)

Serum insulin and insulin like growth factor 1 (IGF1) levels were measured by ELISA using a microplate reader (BioTek Instruments Inc.) and respective kits (Invitrogen) following the manufacturer’s protocol. For the isolation of serum, mouse blood was collected from the tail vein under anesthesia in anticoagulant-free tubes. Blood was then allowed to clot for 30 min at room temperature, and the serum was separated from the clotted blood after centrifugation at 1000 × g for 10 min.

### *In vitro* silencing, blocking and induction of IGF1-IGF1R-mTOR pathway

siRNA for mouse-IGF1R was constructed *in vitro* using the Silencer siRNA Construction Kit (Invitrogen) according to the manufacturer’s protocol. Target-specific and scrambled control siRNA (Sigma-Aldrich) were added (final concentration 50 nM) *in vitro* for 2 h in serum-starved cells in the presence of lipofectamine-2000 (Invitrogen). The primers used are listed below: The siIGF1R was used to block insulin like growth factor 1 receptor (IGF1R) expression. Rapamycin (Merck) (25 pM dissolved in DMSO) was used to block mTORC1 and mouse recombinant IGF1 (R&D Systems) (0.1 μg/ml) was used to induce IGF1R signaling. siIGF1R, rapamycin, and m-rIGF1 were administered alone and in all possible combinations *in vitro* to CD8^+^ T cells isolated from the TDLNs of diabetic and non-diabetic mice.

### *In vitro* antigen stimulation of clinical T1D PBMCs

This study was conducted to determine the functional capabilities of clinical T1D PBMCs. T1D PBMCs (n=7) along with non-diabetic control PBMCs (n=5) were plated in five different 60 mm petriplate (Corning) at a number of 1×10^7^ cells per plate in RPMI-1640 media supplemented with 10% (v/v) heat-inactivated FBS (Himedia). Four group of plate received tumor lysate of MCF-7 (5 μg/ml media), human breast cancer cell line and one group of plate remain as untreated. One group of plates among the four, received rapamycin (25 pM dissolved in DMSO) as mTOR inhibitor, one group of plate received picropodophyllotoxin (PPT) (0.2 μM dissolved in DMSO) as IGF1R inhibitor and another group of plated received both rapamycin and PPT. PBMCs were incubated in a CO_2_ incubator (Thermo Fisher Scientific) at 37 °C with 5% CO_2_ and 95% humidity for 48 hours. After that, cells were stained with fluorescent labelled antibodies for flow cytometry. Culture supernatant was preserved for ELISA.

### Immunofluorescence microscopy

Tumor cryo-sections adhered to glass slides were blocked in 5% BSA (Sisco Research Laboratory) at RT. For intracellular staining, the sections were incubated with 0.15% Triton X-100 prior to blocking. After blocking, the sections were incubated with specific primary antibodies overnight at 4°C followed by FITC and PE-tagged secondary antibodies for 3 h at RT. The sections were then washed and mounted with Fluoroshield and DAPI (Abcam). Images were viewed under a fluorescence microscope (Olympus) with a 40X (NA 1.1, FN 22) objective lens at 24 °C, and images were captured with an attached camera (Olympus) using cellSens standard software (1.18 build 16686, Olympus). The acquired images were analyzed using ImageJ software.

### Quantification and statistical analysis

ImageJ (National Institutes of Health, Bethesda, MD, USA) was used for image analysis, brightness, contrast adjustment, and quantification of the agarose gel bands. Image Lab (Bio-Rad) was used for western blot gel scanning and analysis. The ECAR and OCR data were analyzed using the online Seahorse XF Analyzer software (Agilent). Flow cytometry analyses were done by using FlowJo v10 (BD Bioscience) software. All reported results represent the mean ± SEM of data obtained from two (i*n vivo*; n = 6) or three to six (for *in vitro* assays) independent experiments. Statistical significance was established by unpaired Student’s t-test (for two groups) or two-way analysis of variance followed by Tukey’s post-hoc test (for more than two groups) using the GraphPad Prism 8 software (GraphPad Software, San Diego, CA, USA). For survival analysis, Kaplan-Meier survival analysis followed by data analysis with the log-rank (Mantel-Cox) test was used. Differences between the groups were considered significant at a p-value of 0.05. Search Tool for the Retrieval of Interacting Genes/Proteins (STRING) was used to establish interactome maps of the expressed genes (STRING v.11.5). Graphical representations were created with BioRender.com.

### Data availability

Data generated in this study are available in the article and its supplementary data, or from the corresponding author upon reasonable request.

