## Supplementary Material for "Experimental type 1 diabetes metabolically rejuvenates CD8^+^ T cells for improved control of tumor growth through an IGF1-IGF1R axis"

Supplementary Figure S1

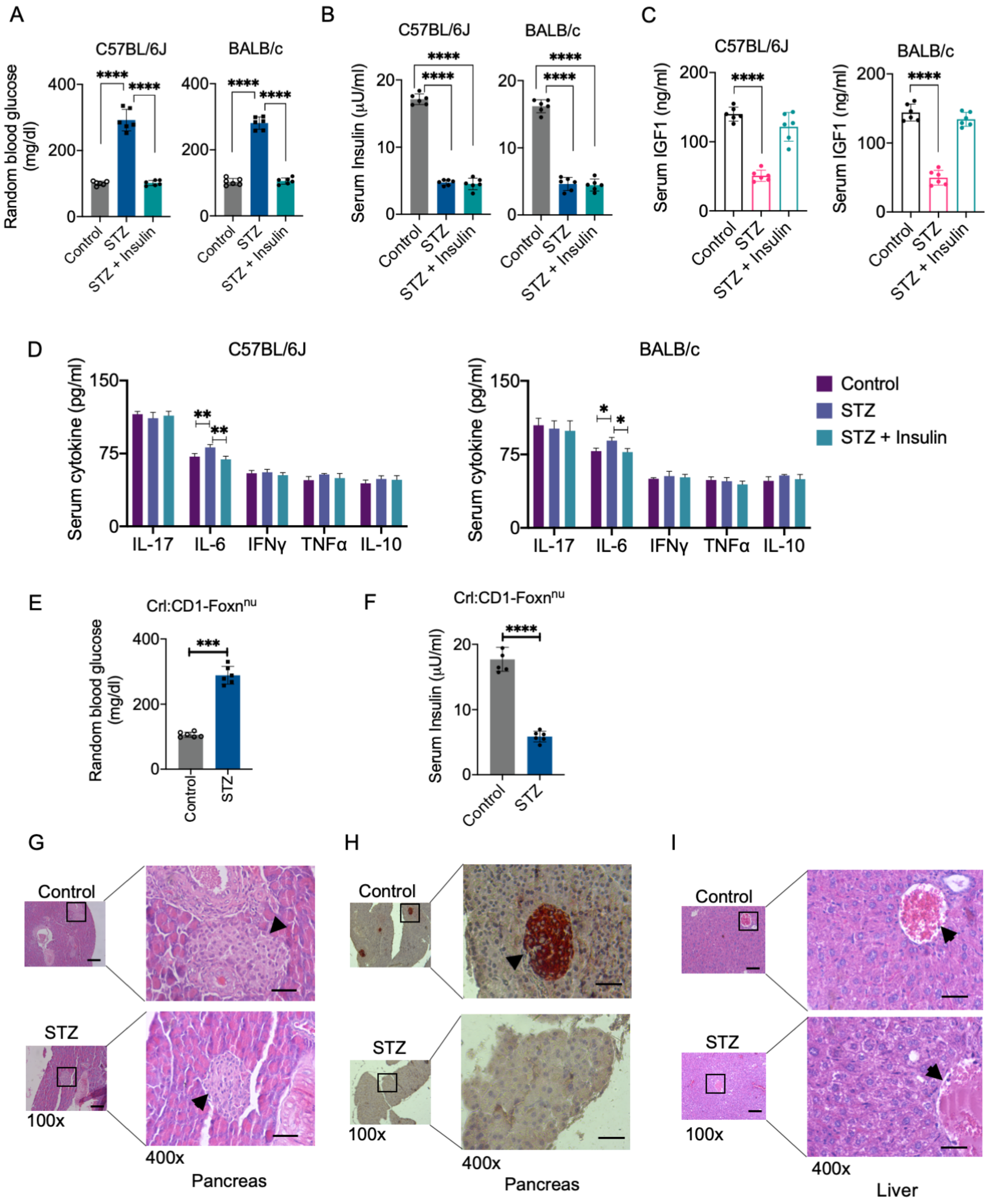


**Supplementary Figure S1.** (A) Random blood glucose level (mg/dl) of C57BL/6J and BALB/c mice from three different groups (n=6) after STZ treatment prior to tumor induction. (B) Serum insulin level (µU/ml) of C57BL/6J and BALB/c and nude mice from three different groups (n=6). Blood was collected prior to that day's insulin injection and before tumor inoculation. (C) Serum insulin like growth factor (IGF1) level (ng/ml) of C57BL/6J and BALB/c from three different groups (n=6) prior to tumor inoculation. (D) Serum cytokine level of IL-17, IL-6, IFNγ, TNFα and IL-10 of C57BL/6J and BALB/c mice from three different groups (n=3) prior to tumor inoculation. (E) Random blood glucose and (F) serum insulin of nude mice (n=6) after STZ treatment before tumor induction. (G) Hematoxylin-eosin stained and (H) immunohistochemistry of pancreas sections with anti-insulin antibody showing reduced pancreatic islets (arrow marked) with STZ treatment from C57BL/6J mice, prior to tumor induction, in 100x and 400x magnification. (I) Hematoxylin-eosin stained untreated and STZ treated liver sections with hepatic portal vein (arrow marked) in 100x and 400x magnification. One way ANOVA for A, B and C, two way ANOVA for D was performed for significance (*P ≤ 0.05, **P ≤ 0.01, ***P ≤ 0.001, ****P ≤ 0.0001)

Supplementary Figure S2.

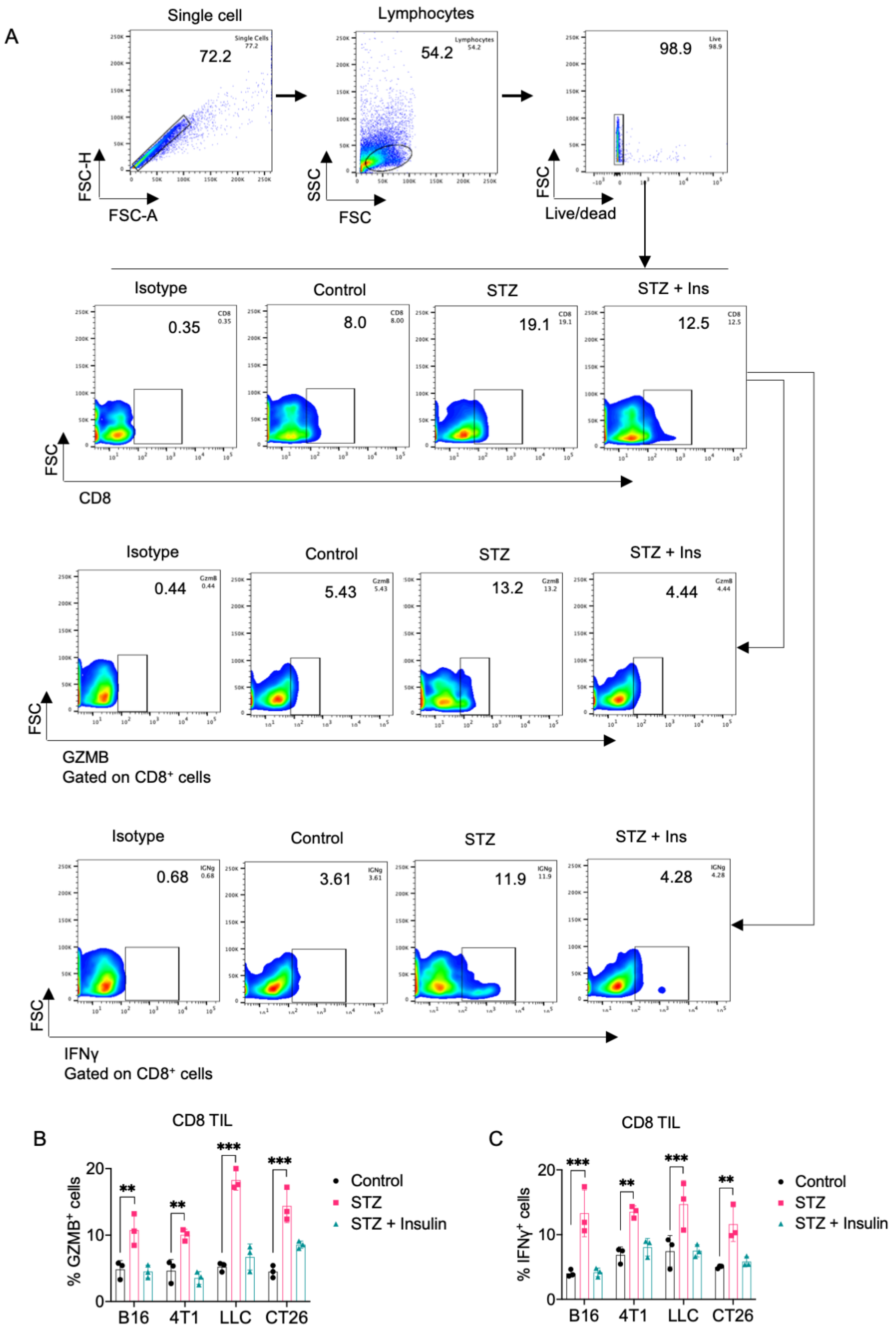


**Supplementary Figure S2.** (A) Gating strategy and representative pseudo color plots of CD8⁺, granzyme B (GZMB⁺) and IFNγ⁺ cells in tumor infiltrating lymphocytes (TIL) of B16F10, 4T1, LLC and CT26 tumors from untreated (Control), STZ treated and STZ+Insulin treated groups. (B) Percent GZMB⁺ and (C) Percent IFNγ⁺ cells in CD8 TIL of B16F10, 4T1, LLC and CT26 tumors from untreated (Control), STZ treated and STZ+Insulin treated groups (n=3). Two way ANOVA for was performed to determine statistical significance. (*P ≤ 0.05, **P ≤ 0.01, ***P ≤ 0.001, ****P ≤ 0.0001)

Supplementary Figure S3.

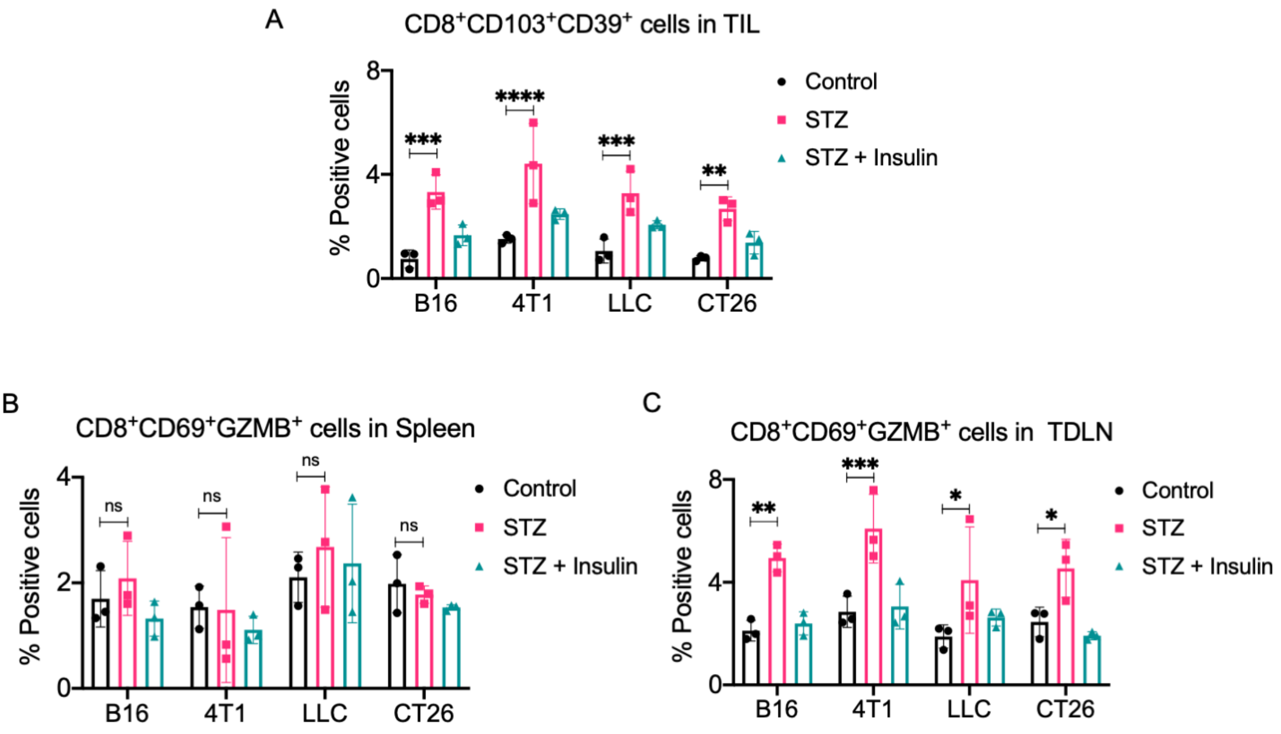


**Supplementary Figure S3.** (A) Percent CD8⁺CD103⁺CD39⁺ cells in tumor infiltrating lymphocytes (TILs) from B16, 4T1, LLC and CT26 tumors of three different groups (n=3). (B) Percent CD8⁺CD69⁺GZMB⁺ cells in spleen from B16, 4T1, LLC and CT26 tumors of three different groups (n=3). (C) Percent CD8⁺CD69⁺GZMB⁺ cells in tumor draining lymph nodes (TDLN) from B16, 4T1, LLC and CT26 tumors of three different groups (n=3). Two way ANOVA was performed to determine statistical significance. (*P ≤ 0.05, **P ≤ 0.01, ***P ≤ 0.001, ****P ≤ 0.0001)

Supplementary Figure S4.

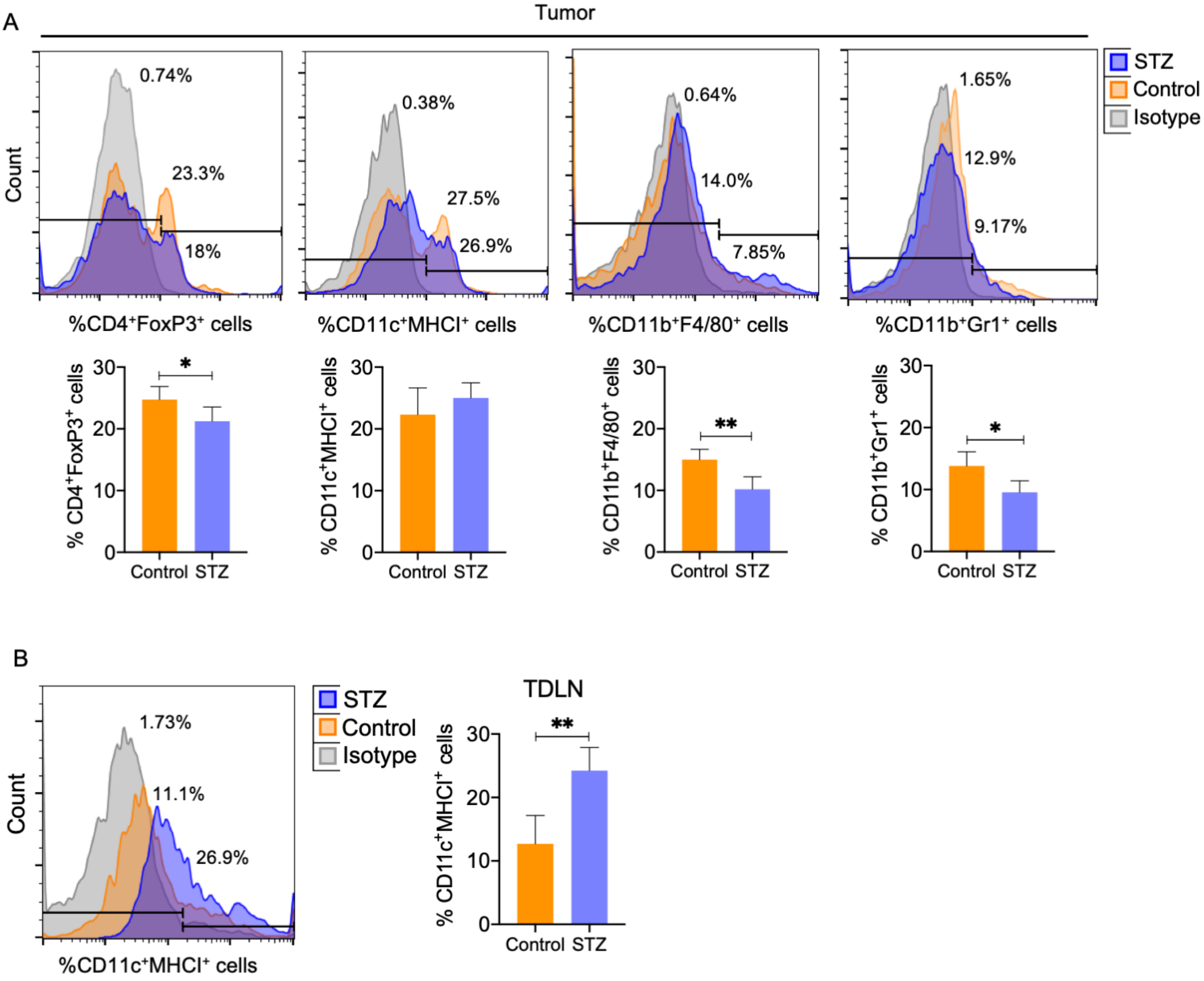


**Supplementary Figure S4.** (A) Histograms and graphs of CD4$^+$FoxP3$^+$ Treg cells, CD11c$^+$MHCI$^+$ dendritic cells (DCs), CD11b$^+$F4/80$^+$ tumor associated macrophages (TAMs) and CD11b$^+$Gr1$^+$ myeloid derived suppressor cells (MDSCs) from tumors of untreated (control) and STZ treated C57BL/6J mice (n=5). (B) CD11c$^+$MHC1$^+$ DC frequency in tumor draining lymph nodes (TDLN) of untreated (control) and STZ treated C57BL/6J mice (n=5). Dendritic cell population in TDLN of STZ and untreated tumor bearing C57BL/6J mice. Unpaired t tests were performed to calculate significance. (*P ≤ 0.05, **P ≤ 0.01, ***P ≤ 0.001, ****P ≤ 0.0001)

Supplementary Figure S5.

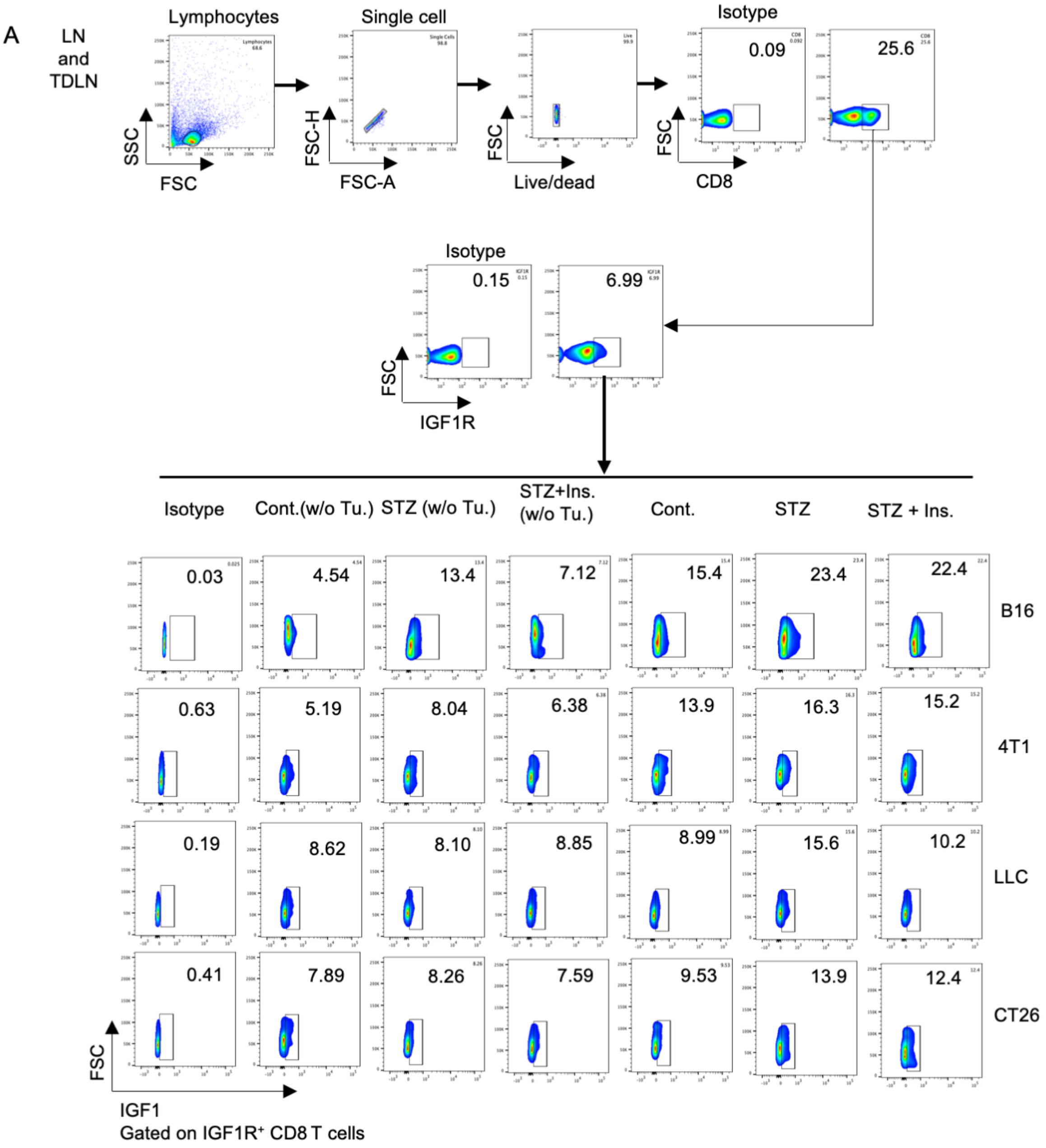


**Supplementary Figure S5.** (A) Flow cytometry gating strategy and representative dot plots of $CD8^{+}IGF1R^{+}IGF1^{+}$ T cells in lymph nodes (LNs) and tumor draining lymph nodes (TDLNs) along with isotype controls. Each experimental set with six groups were analyzed, three without tumor, and three with tumor. Four experimental groups with B16, 4T1, LLC and CT26 tumors were analyzed and graph was presented in Figure 5F.

Supplementary Figure S6.

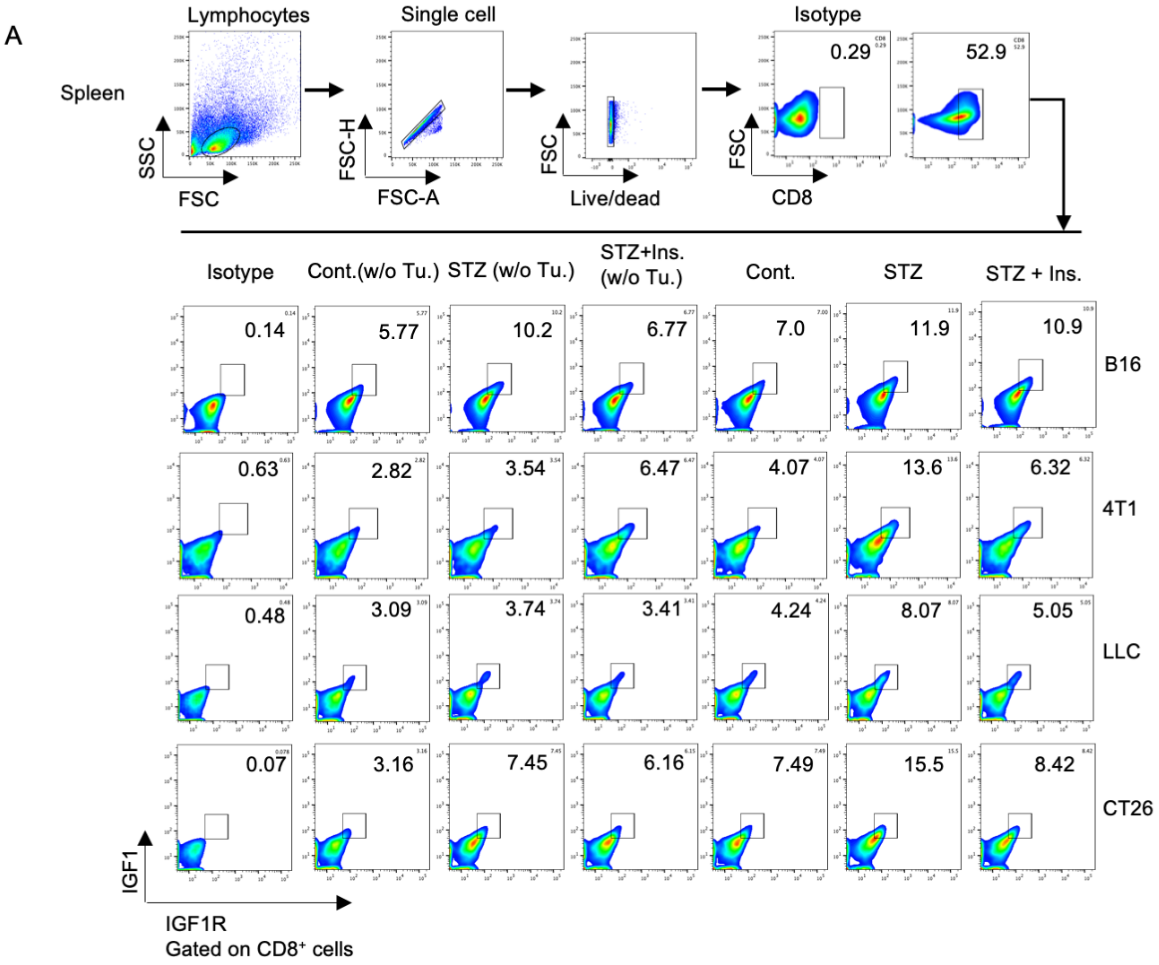


**Supplementary Figure S6.** (A) Flow cytometry gating strategy and representative dot plots of $CD8^+IGF1R^+IGF1^+$ T cells in spleen along with isotype controls. Each experimental set with six groups were analyzed, three without tumor, and three with tumor. Four experimental groups with B16, 4T1, LLC and CT26 tumors were analyzed and graph was presented in Figure 5G.

Supplementary Figure S7.

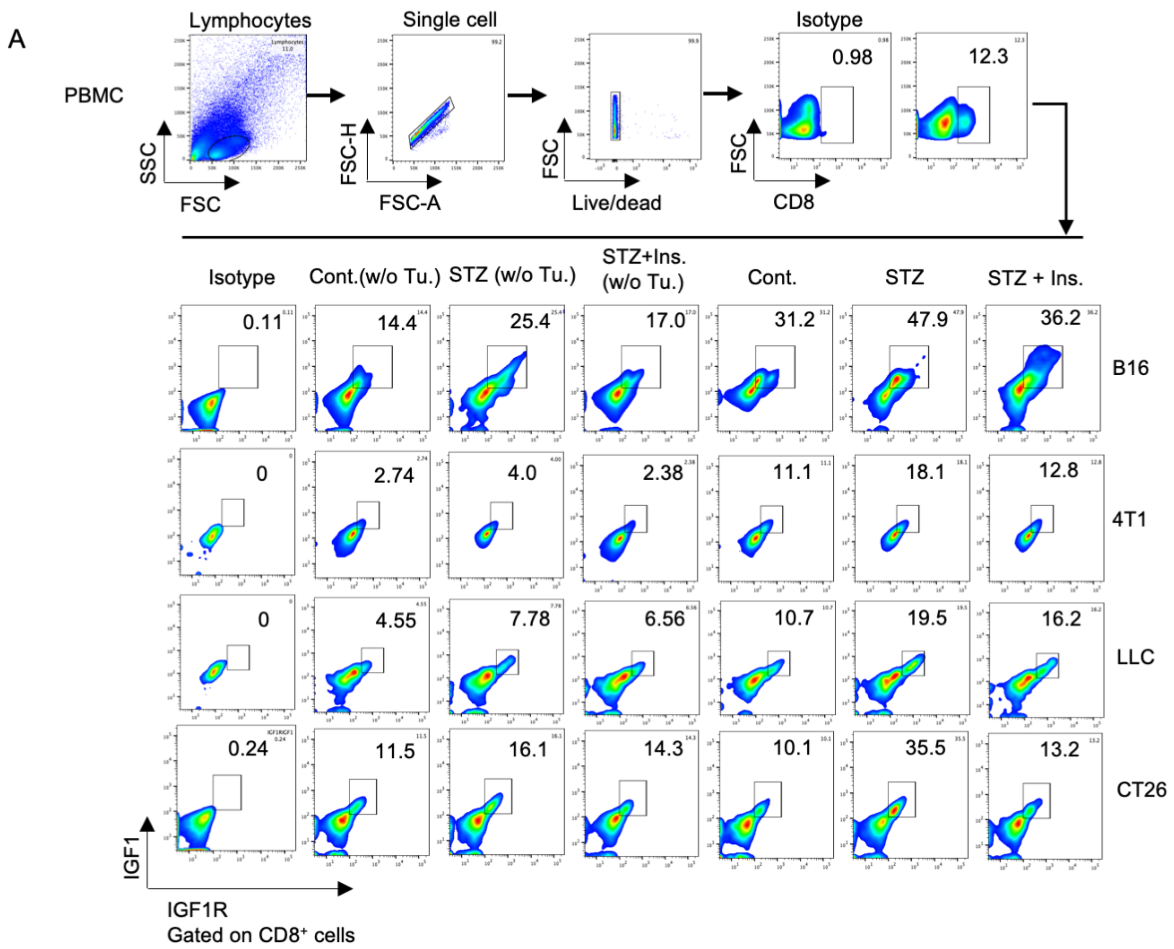


**Supplementary Figure S7.** (A) Flow cytometry gating strategy and representative dot plots of CD8⁺IGF1R⁺IGF1⁺ T cells in peripheral blood mononuclear cells (PBMCs) along with isotype controls. Each experimental set with six groups were analyzed, three without tumor, and three with tumor. Four experimental groups with B16, 4T1, LLC and CT26 tumors were analyzed and graph was presented in Figure 5G.

Supplementary Figure S8

A

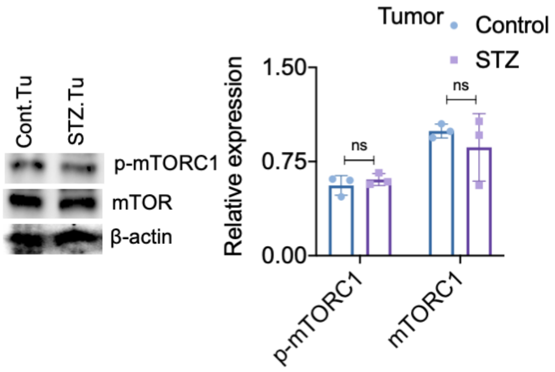


**Supplementary Figure S8.** (A) Western blot of phospho-mTORC1 and mTOR of control and STZ treated B16F10 tumor. Relative expression was measured against β-actin. Two way ANOVA was performed for significance. ns= not significant.

**Supplementary Table ST1.** Type 1 diabetes patient details.

| Sl.No. | Sex | Age | Collected Sample | Glucose Random (mg/dl) | HbA1c | GAD65 antibody | Anti-insulin antibody |
|---|---|---|---|---|---|---|---|
| 1 | F | 18 | Blood | 378 | 7.9 | Positive | Positive |
| 2 | M | 19 | Blood | 213 | 7.6 | Positive | Positive |
| 3 | M | 20 | Blood | 518 | 12.6 | Positive | Positive |
| 4 | F | 23 | Blood | 214 | 8.0 | Positive | Positive |
| 5 | F | 28 | Blood | 229 | 8.7 | Positive | Positive |
| 6 | M | 24 | Blood | 357 | 7.9 | Positive | Positive |
| 7 | M | 24 | Blood | 330 | 8.2 | Positive | Positive |

**Supplementary Table ST2.** Non-diabetic healthy donor details.

| Sl.No. | Sex | Age | Collected Sample | Glucose Random (mg/dl) | HbA1c | GAD65 antibody | Anti-insulin antibody |
|---|---|---|---|---|---|---|---|
| 1 | M | 27 | Blood | 109 | 4.1 | Negative | Negative |
| 2 | M | 25 | Blood | 98 | 5.0 | Negative | Negative |
| 3 | F | 23 | Blood | 95 | 4.6 | Negative | Negative |
| 4 | M | 28 | Blood | 102 | 4.2 | Negative | Negative |
| 5 | F | 26 | Blood | 112 | 5.2 | Negative | Negative |

**Supplementary Table ST3.** Key resources table

| REAGENT or RESOURCE | SOURCE | IDENTIFIER |
|---|---|---|
| **Antibodies** | | |
| PE anti-mouse CD8a | BioLegend | Cat# 100708; RRID:AB_312747 |
| FITC anti-mouse CD8a | BioLegend | Cat# 100705; RRID:AB_312744 |
| APC anti-mouse CD8a | BioLegend | Cat# 100711; RRID:AB_312750 |
| PE-Cy5 Mouse Anti-Human CD8 | BD Pharmingen | Cat#555368 |
| BD IMag Anti-Mouse CD8a Particles - DM | BD Bioscience | Cat# 551516 |
| PE anti-mouse CD28 | BioLegend | Cat# 102106; RRID:AB_312871 |
| FITC anti-mouse CD69 | BioLegend | Cat# 104505; RRID:AB_313108 |
| APC anti-mouse CD183 (CXCR3) | BioLegend | Cat# 155906; RRID:AB_2814078 |
| FITC anti-mouse CD279 (PD-1) | BioLegend | Cat# 135214; RRID:AB_10680238 |
| PE anti-mouse CD223 (LAG-3) | BioLegend | Cat# 125207; RRID:AB_2133344 |

| | | |
|---|---|---|
| PE Mouse Anti-TCF-7/TCF-1 Antibody | BD Biosciences | Cat# 564217; RRID:AB_2687845 |
| Rat Anti-Insulin Monoclonal antibody | R&D Systems | Cat# MAB1417; RRID:AB_2126533 |
| IGF1 Monoclonal Antibody | Thermo Fisher Scientific | Cat# MA5-23772; RRID:AB_2609723 |
| Mouse Anti-Human IGF1R Monoclonal antibody | R&D Systems | Cat# MAB391; RRID:AB_2122409 |
| Anti-β-Actin Antibody | Sigma-Aldrich | Cat# A5441; RRID:AB_476744 |
| Anti-mTOR Antibody | Cell Signaling Technology | Cat# 2972; RRID:AB_330978 |
| Anti-phospho-mTOR | Cell Signaling Technology | Cat# 2971; RRID:AB_330970 |
| FITC anti-mouse CD4 | BioLegend | Cat# 100406; RRID:AB_312691 |
| PE anti-mouse FOXP3 | BioLegend | Cat# 118904; RRID:AB_2936574 |
| FITC anti-mouse CD11c | BioLegend | Cat# 117306; RRID:AB_313775 |
| PE anti-mouse/human CD11b | BioLegend | Cat# 101208; RRID:AB_312791 |
| FITC anti-mouse F4/80 Recombinant | BioLegend | Cat# 157309; RRID:AB_2876535 |
| FITC anti-mouse Ly-6G/Ly-6C (Gr-1) | BioLegend | Cat# 108405; RRID:AB_313370 |
| Goat anti-Mouse IgG (H+L) Secondary Antibody, HRP | Thermo Fisher Scientific | Cat# 31430; RRID:AB_228307 |
| Mouse Anti-Hamster IgG PE MAb | R&D Systems | Cat# F0120; RRID:AB_1856861 |
| Goat anti-Mouse IgG (H+L) Secondary Antibody, FITC | Novus Biologicals | Cat# NB720-F; |
| **Biological samples** | | |
| Type-1 diabetic patient blood | Doyen Diagnostics and Research Foundation, Kolkata, India | NA |
| **Chemicals, peptides, and recombinant proteins** | | |
| Streptozotocin | Millipore | Cat# 572201 |
| DMEM high glucose | Himedia | Cat# AL007, |
| FBS heat inactivated | Himedia | Cat# RM10975, |
| Collagenase | Merck | Cat# C4-BIOC, |
| Lymphocyte separation medium, human | Himedia | Cat# HiSep LSM 1077, |
| Lymphocyte separation medium, mouse | Himedia | Cat# HiSep LSM 1084, |

| | | |
|---|---|---|
| Trizol RNA extraction | Invitrogen | Cat# 15596018, |
| 2x green master mix | Promega | Cat# M7122, |
| Cytofix/Cytoperm buffer | BD Bioscience | Cat# BDB554714 |
| Lipofectamine-2000 | Invitrogen | Cat# 11668019 |
| Rapamycin (mTOR inhibitor) | Merck | Cat# R8781, |
| Mouse recombinant IGF1 | R&D Systems | Cat# 791-MG-050, |
| Picropodophyllotoxin (IGF1R inhibitor) | Sigma-Aldrich | Cat# T9576-1MG |
| **Critical commercial assays** | | |
| LDH Cytotoxicity Assay Kit | Roche | Cat# 4744926001 |
| Seahorse XF Glycolytic Rate Assay Kit | Agilent | Cat# 103344-100, |
| First Strand cDNA Synthesis Kit | Thermo Scientific | Cat# K1622, |
| ECL Western Blotting Substrate Kit | Advansta | Cat# K-12045-D50 |
| Insulin ELISA kit | Invitrogen | Cat# EMINS |
| IGF1 ELISA kit | Invitrogen | Cat# EMIGF1 |
| Silencer siRNA Construction Kit | Invitrogen | Cat# AM1620, |
| **Experimental models: Cell lines** | | |
| B16F10 | NCCS, Pune, India | RRID:CVCL_XH27 |
| 4T1 | NCCS, Pune, India | RRID:CVCL_0125 |
| LLC | NCCS, Pune, India | RRID:CVCL_3009 |
| CT-26 | NCCS, Pune, India | RRID:CVCL_7254 |
| **Experimental models: Organisms/strains** | | |
| C57BL/6J | Jackson Laboratories, USA | RRID:IMSR_JAX:000664 |
| BALB/c | State Centre for Laboratory Animal Breeding, West Bengal Livestock Development Corporation Limited, India | |
| Crl:CD1-Foxn1nu | CDRI, Lucknow | RRID:IMSR_CRL:086 |
| **Oligonucleotides** | | |
| siIGF1R sense: 5' AACCCTCCTCCGGAGCCAGACCCTGTCTC 3' siIGF1R antisense: 5' AAGTCTGGCTCCGGAGGAGGGCCTGTCTC 3' | Integrated DNA Technologies | |
| **Software and algorithms** | | |
| ImageJ | Fiji | |
| FlowJo 10.8.1 | BD Biosciences | |
| Seahorse XF Analyzer software | Agilent | |
| GraphPad Prism 8 | GraphPad Software Inc. | |
| STRING v.11.5 | Global Core Biodata Resource | |
| BioRender | Biorender Inc. | |

**Supplementary Table ST4.** Primer sequences used to study mRNA expressions of genes by RT-PCR along with their melting temperatures ($T_m$) and primer lengths.

| Name of gene | Primers 5'-3' | $T_m$ (°C) | Primer length (base) |
|---|---|---|---|
| *perforin* | Forward: GATGTGAACCCTAGGCCAGA<br>Reverse: GGTTTTTGTACCAGGCGAAA | 59<br>55 | 20<br>20 |
| *granzyme b* | Forward: TCGACCCTACATGGCCTTAC<br>Reverse: TGGGGAATGCATTTTACCAT | 59<br>53 | 20<br>20 |
| *ifnγ* | Forward: GGTGACATGAAAATCCTGCAGAGC<br>Reverse: TCAGCAGCGAGTGATTTTCCGCTT | 58<br>61 | 24<br>24 |
| *il-2* | Forward: GCAGGCCACAGAATTGAAAG<br>Reverse: TCCACCACAGTTGCTGACTC | 57<br>59 | 20<br>20 |
| *β-actin* | Forward: CAACCGTGAAAAGATGACCC<br>Reverse: ATGAGGTAGTCTGTCAGGTC | 57<br>57 | 20<br>20 |
| *glut 1* | Forward: CAGGTGTTTGGCTTAGACTC<br>Reverse: GGATCAGATGCAAAGCTTTC | 57<br>57 | 20<br>20 |
| *g-6-pd* | Forward: GGCCAGTTTCTATGAGGAGTATG<br>Reverse: GCTGATGTTGAGAGGCAGTT | 65<br>65 | 23<br>20 |
| *pkm 2* | Forward: ACTGGCATCATTTGTACCAT<br>Reverse: GGATCAGATGCAAAGCTTTC | 54<br>55 | 20<br>20 |
| *ldh a* | Forward: CCTGAAGTCTCTTAACCCAG<br>Reverse: CCGCCTAAGGTTCTTCATTA | 54<br>54 | 20<br>20 |
| *pcx 1* | Forward: GTAAAGACCAACATCCCCTT<br>Reverse: ATGACATGTCCGAGGTAATG | 54<br>54 | 20<br>20 |
| *pdk 1* | Forward: CTCCTTATTGTTCGGTGGAA<br>Reverse: TTGTATTGTCTGTCCTGGTG | 55<br>54 | 20<br>20 |
| *pdk 3* | Forward: AACTCTCGTTACTCTGGGTA<br>Reverse: TACAGACGAGAAATTGGCAA | 54<br>55 | 20<br>20 |
| *idh 1* | Forward: GTGGAGATGCAAGGAGATGAA<br>Reverse: ATGCAGATCCAGTTCCACATAG | 64<br>64 | 21<br>22 |
| *idh 2* | Forward: AACACCGACGAGTCCATTTC<br>Reverse: TCAAGTAGAGCGGCCATTTC | 65<br>65 | 20<br>20 |
| *fh 1* | Forward: TGCTGAAGTAAACCAGGAGTATG<br>Reverse: CCAAACCACCAGAGGAAAGT | 64<br>65 | 23<br>20 |
| *igf 1* | Forward: TCTGAGGAGACTGGAGATGT<br>Reverse: TAGGTCTTGTTTCCTGCACT | 54<br>55 | 20<br>20 |
| *igf 1R* | Forward: GAGATGACCAATCTCAAGGA<br>Reverse: CTTGTTCCCCACAATGTAGT | 55<br>54 | 20<br>20 |
| *igf 2* | Forward: GGCAAGTTCTTCCAATATGA<br>Reverse: CTTTGGGTGGTAACACGAT | 55<br>55 | 20<br>19 |
| *insulin R* | Forward: ATACCATGAATTCCAGCAAC<br>Reverse: ATGTTGATGATCAGGCTACC | 54<br>54 | 20<br>20 |
